# Anti-defense systems of viral communities vary with genomic traits, host immunity and environmental stress

**DOI:** 10.64898/2026.09.28.755093

**Authors:** Dan Huang, Yiyan Yang, Ruonan Wu, Jose Luis Balcazar, Pedro J.J. Alvarez, Jizhong Zhou, Pingfeng Yu

**Affiliations:** State Key Laboratory of Soil Pollution Control and Safety, College of Environmental and Resource Sciences, Zhejiang University, Hangzhou 310058, China; Channing Division of Network Medicine, Department of Medicine, Brigham and Women’s Hospital and Harvard Medical School, Boston, MA 02115, USA; Earth and Biological Sciences Directorate, Pacific Northwest National Lab, Richland, WA, 99352, USA; Catalan Institute for Water Research (ICRA-CERCA), Girona, 17003, Spain; Department of Civil and Environmental Engineering and Rice WaTER Institute, Rice University, Houston, Texas 77005, USA; Institute for Environmental Genomics, University of Oklahoma, Norman, OK 73019, USA

**Keywords:** Viral anti-defense systems, virus-host interaction, viral ecology, heavy metal resistance, environmental stress

## Abstract

**Background:** Despite the ecological importance of viral anti-defense systems (ADS), yet how viral ADS repertoires vary across microbial ecosystems remains poorly understood. Here, we integrated global virus datasets with metagenomes from heavy metal-contaminated soils and chromium-stress microcosms to investigate ecological determinants of viral ADS.

**Results:** We found that ADS repertoires were constrained by viral genome capacity, with most ADS-carrying viruses encoded a single system, whereas multi-ADS occurred mainly in larger genomes. Viral lifestyle partitioned ADS, with temperate viruses enriched in ADS targeting early-stage defenses and virulent viruses preferentially encoding ADS against mid- and late-stage defense. ADS composition also varied by host lineage and across habitats, linking viral ADS repertoires to host immune landscapes and ecosystem-specific viral strategies. Moreover, along the heavy metal contamination gradient, changes in the prevalence of ADS-carrying viruses occurred alongside shifts in prokaryotic defense profiles. In addition, expanded ADS repertoires may further increase dissemination potential of MGE-borne adaptive genes, highlighting viral ADS as a potential contributor to the adaptive evolution of microbial community.

**Conclusions:** These findings reveal that viral ADS are shaped by genomic traits, host immunity and environmental stress, with implications for virus-host interaction and microbial adaptive evolution. More broadly, this study extends anti-defense research beyond individual virus-host systems, offering an integrative, community-level perspective to guide future studies testing whether and how virus-prokaryote immune interactions contribute to microbial community dynamics and evolution in changing environments.

## Introduction

Microbial ecosystems are continually shaped by evolution between prokaryotes and their viruses[1–3]. These interactions influence not only microbial mortality and community composition, but also horizontal gene transfer and biogeochemical cycling across ecosystems[4–6]. To resist viral infection, prokaryotes have evolved a remarkable diversity of antiviral defense systems[7–11], including CRISPR-Cas, restriction-modification (RM) systems, and many newly discovered systems[12–15]. In response, viruses have evolved diverse strategies to counter host immunity[16, 17], ranging from genetic variations that enables immune evasion to dedicated anti-defense systems (ADS) that suppress, disrupt, or bypass host defense mechanisms[18–20]. As major viral counter-defense mechanisms, ADS represent adaptive responses to defense-mediated selection pressures and play a central role in shaping virus-host interaction dynamics[16]. Furthermore, because many prokaryotic defense systems restrict not only viruses but also plasmids and other mobile genetic elements (MGEs), ADS-like mechanisms have been identified across diverse MGEs[21–23]. Consequently, the ecological significance of viral ADS may extend beyond infection outcomes, with the potential to influence horizontal gene transfer, MGE dissemination, and the spread of adaptive traits across microbial communities[21, 24]. Despite the ecological importance of ADS, the determinants of ADS repertoire composition in viral communities and the consequences of viral ADS for microbial adaptation across complex ecosystems remain unclear.

Previous ADS studies have largely focused on discovering new mechanisms that suppress or bypass specific host defense systems, often using single virus-host model systems[20, 25–27]. Although this mechanism-centered perspective has greatly expanded our understanding of the diversity of viral counter-defense strategies, it has not yet provided an integrative ecological framework for explaining how ADS repertoires are organized across complex microbial communities. Viral populations differ substantially in genome capacity and infection lifestyle[28, 29]. Genome capacity may limit the number of ADS that can be encoded and maintained alongside essential replication functions[29], whereas alternative infection strategies expose viruses to different immune barriers and may therefore favor distinct ADS repertoires. Prokaryotic host lineages also vary markedly in their immune architectures, generating different selective landscapes that can shape the maintenance and diversification of ADS[7, 30]. More broadly, environmental conditions may influence ADS repertoires by altering microbial community composition, host physiological states, viral lifestyles, and the intensity of virus-host interactions[31–33]. We therefore propose that ADS repertoires may be shaped by ecological filtering operating simultaneously across viral traits, prokaryotic host immunity, and environmental conditions.

Here, we integrated global viral genome collections from IMG/VR[34] and VIRE[35], large-scale metagenomes from 401 heavy metal-contaminated soils across China, and chromium-stress microcosm experiments to systematically investigate the composition, determinants, and potential ecological consequences of viral ADS in microbial communities (**Fig. 1**). First, using global high-quality viral genomes, we examined whether viral genome size, infection lifestyle, prokaryotic host, and habitat type jointly shape ADS repertoires across viral communities. We then used heavy metal-contaminated soil metagenomes to validate these global-scale patterns and further assess whether environmental stress alters virus-host immune interactions and the distribution of ADS-carrying viruses. To experimentally validate the effect of environmental stress on viral ADS, we established chromium-stress microcosms to characterize ADS-carrying viruses under different chromium stress levels. Finally, to assess the broader ecological impacts of viral ADS, we used anti_CRISPR systems as a model to examine their associations with mobile genetic elements and the dissemination of adaptive resistance genes in heavy metal-contaminated soils. Overall, this study moves ADS research beyond the molecular mechanisms of individual systems toward the ecological determinants and functional consequences of ADS repertoires in complex microbial communities, advancing our understanding of viral counter-immunity, virus-host interactions, and microbial adaptation under environmental stress.

**Fig. 1.**
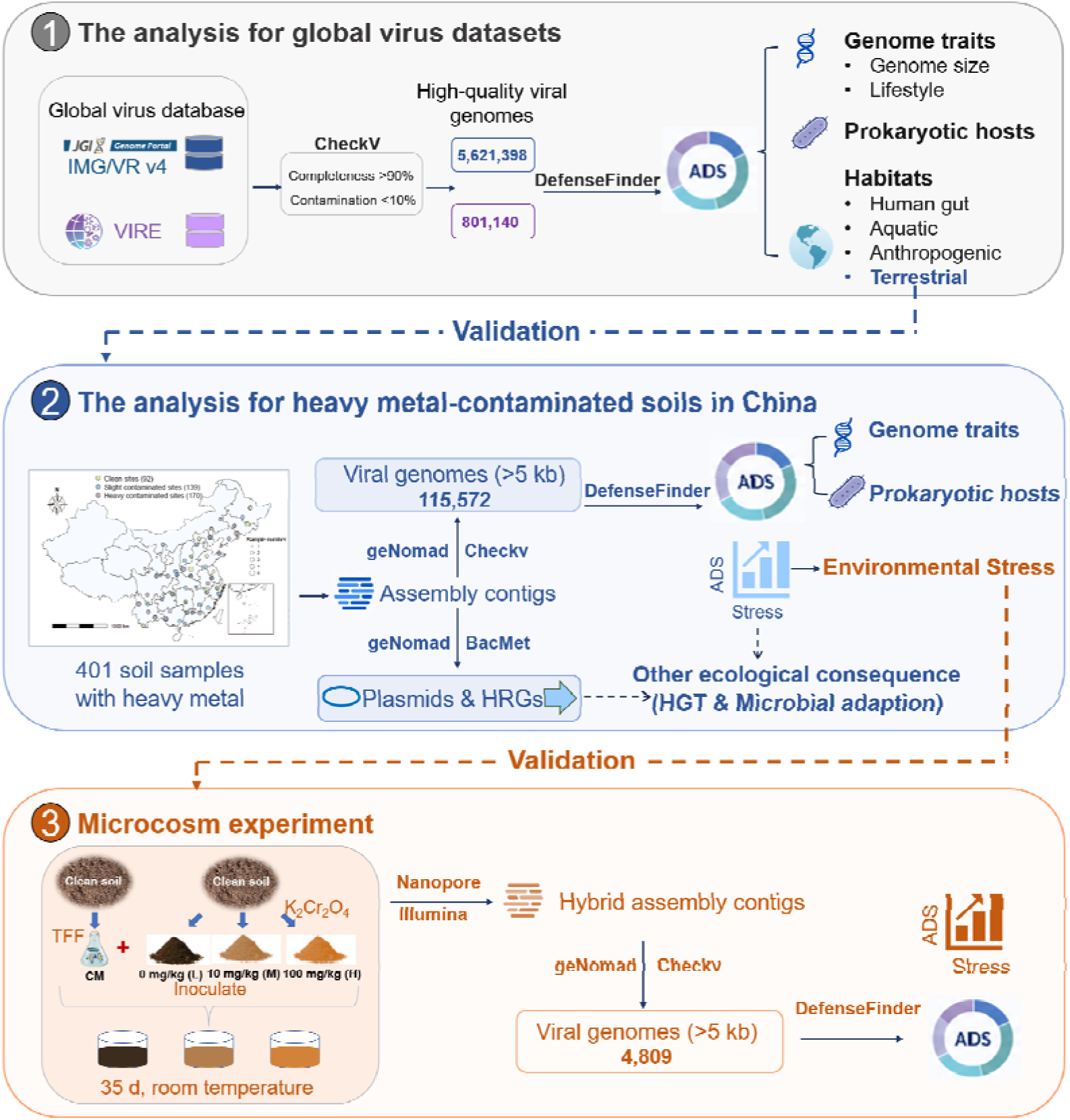
Overview of study design and analytical workflow. Global viral genome collections from IMG/VR v4 and VIRE were filtered using CheckV to obtain high-quality viral genomes with completeness >90% and contamination <10%, followed by annotation of ADS using DefenseFinder[37]. These datasets were used to characterize viral ADS repertoires and evaluate the effects of viral genome traits, including genome size and infection lifestyle, as well as prokaryotic host lineage and habitat type, to ADS repertoire composition. Large-scale metagenomes from 401 heavy metal-contaminated soils across China were then used to validate the global-scale patterns and assess how environmental stress shapes ADS-carrying viral communities and evaluate their potential ecological consequences for microbial adaptation. Chromium (VI)-stress microcosm experiments provided further experimental validation of ADS responses under controlled stress conditions, using hybrid assemblies of short- and long-read sequencing data to recover viral genomes and quantify ADS patterns across chromium (VI) treatments

## Methods

### Global virus dataset from IMG/VR and VIRE database

Two global viral genome resources, IMG/VR[34] and VIRE[35], were used in this study for large-scale analyses of viral ADS. Viral genomes from the IMG/VR database (accessed in March 2025) were first processed using CheckV (v1.0.1), and only sequences with estimated completeness >90% and contamination <10% were retained for downstream analyses, resulting in 5,621,398 high-quality viral genomes. In addition, viral genomes representing terrestrial, aquatic, human gut, and anthropogenic environments were retrieved from the VIRE database[35]. The same completeness and contamination filtering criterion was applied to the VIRE dataset, resulting in 801,140 high-quality viral genomes. Only 3.1% of high-quality VIRE genomes showed high similarity to IMG/VR sequences, based on a threshold of >95% sequence similarity and >80% alignment coverage, indicating limited overlap between the two datasets. To minimize database-specific biases, IMG/VR and VIRE datasets were analyzed independently using the same downstream workflow. All retained viral genomes were further subjected to viral lifestyle classification using PhaTYP[36] and ADS identification using DefenseFinder (v3.0.0, -A --antidefensefinder-only)[37].

### Large-scale heavy metal-stressed soil metagenomic data collection

We compiled a large-scale soil metagenomic dataset by retrieving publicly available shotgun sequencing data from five projects in the NCBI BioProject database (PRJNA1253350, PRJNA1253357, PRJNA1253358, PRJNA1253360 and PRJNA1254387). These studies investigated heavy metal contamination across diverse soil environments in China and included associated physicochemical measurements[38]. The final dataset comprised 401 soil samples spanning a broad gradient of heavy metal contamination (**Fig. 1**). Detailed physicochemical properties of all samples are provided in **Table S1**. The primary heavy metal contaminants in these samples are Pb, Zn, Cu, As and Fe.

### Heavy metal pollution index and sample grouping

To quantify the combined effects of multiple heavy metals, pollution intensity was evaluated using the Nemerow pollution index (P ), calculated based on seven metals (Cd, As, Pb, Cr, Cu, Ni and Zn). Threshold values (S ) were defined according to the Chinese soil environmental quality standard (GB 15618-2018) and adjusted based on soil pH to account for metal bioavailability.

For each metal, a single-factor pollution index was calculated as

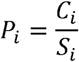

where C represents the measured concentration. The overall pollution index was then computed as:

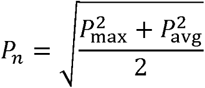

where Pmax is the maximum P and P_avg_ is the mean across detected metals. Only measured metals were included for each sample to avoid underestimation.

Samples were grouped into three categories based on P values: clean (P ≤ 0.7, n = 92), slightly contaminated (0.7 < P ≤ 10.0, n = 139), and heavily contaminated (P > 10.0, n = 170). These categories were used to define a relative ecological gradient for comparative analyses rather than formal environmental risk classification.

### Metagenomic processing and assembly

Raw metagenomic reads were processed using fastp (v0.23.2) to remove adapter sequences and low-quality bases[39]. High-quality reads were assembled using MEGAHIT (v1.2.9) with default parameters[40], with k-mer sizes ranging from 21 to 141 and a minimum contig length of 500 bp. Assembly quality was evaluated using QUAST (v5.2.0)[41]. To enable downstream analyses, contigs shorter than 500 bp were excluded unless otherwise specified. Clean reads were also taxonomically profiled using Kraken2 to obtain an overview of microbial community composition[42].

### Identification of heavy metal resistance genes and normalization

HRGs were identified using the ARGs-OAP pipeline[43], with the default reference database replaced by the experimentally validated BacMet database. Clean reads were mapped to the BacMet database to identify HRG-like sequences[44]. To enable normalization at the cellular level, bacterial cell abundance was estimated using ARGs-OAP based on single-copy marker genes. The abundance of HRGs and other functional features was then normalized per cell to allow cross-sample comparisons.

### Viral identification, classification and host prediction

Viral and plasmid sequences were identified from assembled contigs using geNomad (v2.1.1) in end-to-end mode[45]. To ensure sequence quality, viral contigs shorter than 5 kb were excluded, and potential host contamination and redundancy were removed using CheckV[46]. Taxonomic classification of viral sequences was performed using PhaGCN. [47]. Viral lifestyle (temperate or virulent) was predicted using PhaTYP[36]. Host prediction was conducted using iPHOP, which integrates multiple complementary approaches to improve prediction accuracy[48]. Only high-confidence virus-host associations were retained for downstream analyses. The relative abundance of viral contigs was estimated using CoverM based on read mapping (-m TPM, --trim-min 0.10, --trim-max 0.90,--min-read-percent-identity 0.95, --min-read-aligned-percent 0.80)[49].

### Identification of defense and anti-defense systems

Prokaryotic antiviral defense systems and viral ADS were identified using DefenseFinder (v1.3.0) with default parameters[7, 37]. Input data included assembled contigs from metagenomes, MAGs, plasmids, and viral genomes. To ensure annotation reliability, only complete defense and anti-defense systems with all required genes co-localized on the same genomic fragment were retained[7].

### Microcosm experiment and virome analysis

Soil for microcosm experiments was collected from a relatively clean site adjacent to a chromium-contaminated area in Xining, China, and soil physicochemical properties were measured (**Table S2**). Microbial inoculation was prepared from clean soil using buffer extraction and filtration. Sterilized soils were amended with Cr(VI) to final concentrations of 0, 10, and 100 mg/kg to represent increasing metal stress, and inoculated to establish three treatments (C-0, C-10, and C-100). Microcosms were incubated at 25 °C for 35 days under controlled moisture conditions. Additional experimental details are provided in our previous study[50].

Genomic DNA was extracted from microcosm soils using the Qiagen DNeasy PowerSoil Kit. All samples were subjected to both Illumina NovaSeq 6000 short-read sequencing and Oxford Nanopore long-read sequencing, generating approximately 20 Gb and 3 Gb of data per sample, respectively, as previously described[51]. Briefly, Illumina reads were quality filtered using fastp (v0.23.2), Nanopore reads were processed for adapter removal and quality assessment, and hybrid assemblies were generated using OPERA-MS(v0.9.0)[52]. We deposited the raw sequencing data in the NCBI Sequence Read Archive under accession number PRJNA1104006. Open reading frames were predicted from assembled sequences and clustered to construct a non-redundant gene catalog. Gene abundance was quantified by mapping reads to the gene catalog and normalizing as RPKM. Functional annotation was performed against the KEGG and eggNOG databases[53], and heavy metal resistance genes were identified using the BacMet database[54]. MAG reconstruction, taxonomic annotation, viral identification, and identification of defense systems and ADS were performed as described above.

For host prediction, viral sequences from microcosm datasets were assigned to potential hosts using iPHoP[48] with locally reconstructed MAGs as reference genomes. To improve prediction accuracy, MAGs obtained from each microcosm were incorporated into the host database together with their GTDB-based taxonomic annotations. To ensure reliability, only virus-host associations in which both the viral sequence and its predicted host MAG originated from the same treatment were retained for downstream analyses.

### Causal DAG framework and total effect estimation of ADS on heavy metal resistance

A directed acyclic graph (DAG) was constructed to specify the assumed relationships among variables and to identify the adjustment set for total effect estimation[55]. anti_CRISPR relative abundance was defined as the exposure, and total HRG abundance as the outcome. The Nemerow integrated pollution index and CRISPR-Cas system abundance were included as measured confounders in the adjustment set. Plasmid abundance and temperate virus proportion were not included in the total effect model to avoid adjustment for potential mediators. All continuous variables were normalized by total abundance and transformed using log(x + 1). Variables were subsequently standardized prior to regression analysis. The total effect of anti_CRISPR on HRG abundance was estimated using multivariable linear regression. Inverse probability weighting (IPW) was applied to construct a weighted pseudo-population [56]. Samples were categorized into high and low anti_CRISPR groups, and the probability of assignment to the high anti_CRISPR group was estimated using a generalized linear model. Stabilized weights were calculated and incorporated into weighted regression models. Covariate balance was assessed using standardized mean differences (SMD), with |SMD| < 0.1 considered indicative of adequate balance[57]. A doubly robust model was implemented by including both stabilized weights and covariates in the regression model[58].

### Mediation analysis

A parallel mediation analysis framework was applied with anti_CRISPR abundance as the exposure and total HRG abundance as the outcome[59]. Temperate virus proportion and plasmid abundance were included as two parallel mediators. The residual direct effect was retained in the model. Mediator models were constructed to estimate the associations between anti_CRISPR and each mediator. An outcome model was fitted including the exposure, mediators, and confounders to estimate the associations between mediators and HRG abundance. Indirect effects were estimated using bootstrap resampling, and 95% confidence intervals were calculated[60]. The relative contribution of each mediation pathway was calculated as the ratio of the indirect effect to the total effect (indirect effect / total effect × 100%).

### Statistical analysis

All statistical analyses were performed using Python (version 3.9) and R (version 4.3.1). Pairwise comparisons between groups were conducted using the Wilcoxon rank-sum test. Correlation analyses were performed using Pearson correlation coefficients.

## Results and discussion

### Viral genome size constrains the capacity of encoding anti-defense systems

To investigate the global ecological patterns of viral ADS, we analyzed viral genome collections from the IMG/VR and VIRE databases. The final datasets comprised 5,621,398 and 801,140 high-quality viral genomes (i.e., completeness >90% and contamination <10%) from IMG/VR and VIRE, respectively. ADS were identified in 92,211 IMG/VR viral genomes and 54,465 VIRE viral genomes, representing 101,594 and 59,801 ADS and spanning 14 and 13 major ADS classes, 139 and 99 subtypes, respectively. Among the detected ADS, anti_RM, anti_Thoeris, and anti_CRISPR represented the dominant categories in both datasets, collectively accounting for more than 70% of all identified systems (**Fig. 2a, Supplementary Text S1**). Additional ADS classes, including anti_RecBCD, anti_CBASS, NADP, and anti_Dnd, were also recurrently detected across datasets.

**Fig. 2.**
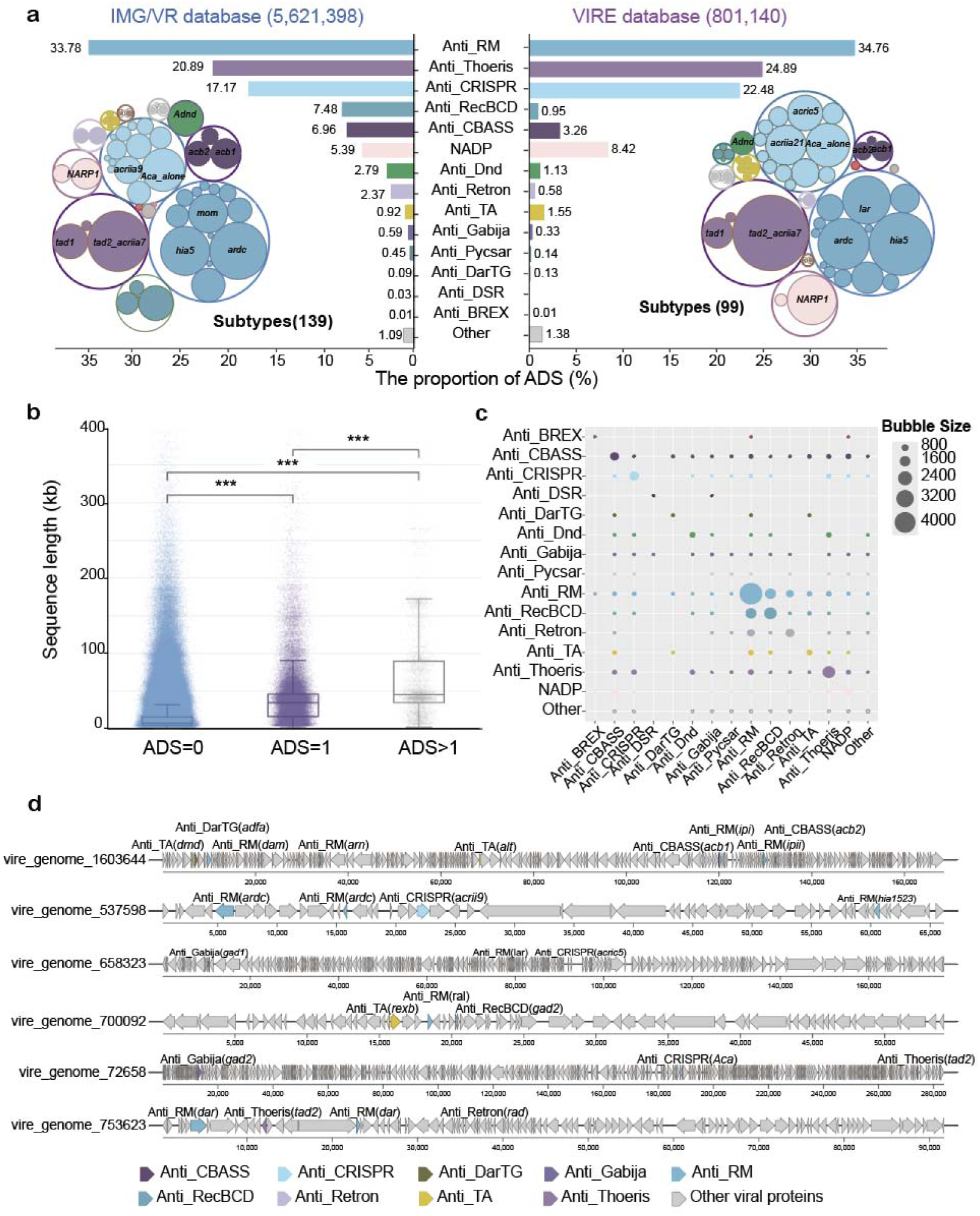
Global distribution patterns and genome size associations of viral ADS. (**a**) Composition of ADS identified in high-quality viral genomes from the IMG/VR (left) and VIRE (right) databases. Viral genomes were filtered using CheckV, retaining only sequences with estimated completeness >90% and contamination <10%. A total of 5,621,398 IMG/VR viral genomes and 801,140 VIRE viral genomes were included in the analyses. ADS were identified using DefenseFinder[37], and the relative abundance of each ADS category was calculated as the proportion of total detected ADS within each dataset. (**b**) Genome size distributions of viral genomes carrying different numbers of ADS in IMG/VRdatabase. Viral genomes were grouped according to the number of encoded ADS (0, 1 or ≥2 ADS). Boxplots show the distribution of viral genome lengths within each category. Center lines indicate medians, boxes indicate interquartile ranges (IQRs), and whiskers represent 1.5 × IQR. Statistical significance was assessed using two-sided Wilcoxon rank-sum tests (***, *P* < 0.001). (**c**) Pairwise co-occurrence frequencies of ADS within the same viral genomes in the IMG/VR database. Bubble sizes indicate the number of viral genomes carrying each ADS combination, highlighting preferential co-occurrence patterns among major ADS classes. (**d**) Representative genome architecture of virus encoding multiple ADS.

Unlike prokaryotic genomes that often encode multiple antiviral defense systems[61], most ADS-carrying viruses (>92%) contained only a single detectable ADS, whereas viruses with multiple detected ADS were comparatively rare. If all defense systems encoded by a host were equally effective against a given virus, successful infection might require that virus to encode multiple corresponding ADS. The predominance of a single detectable ADS instead raises the possibility that only a subset of the host defense repertoire is effective against a particular virus and that virus-specific antiviral activity may extend beyond CRISPR–Cas systems. For example, the GmrSD type IV R-M system selectively targets viruses such as T4[62],whereas Cas9 does not[63]. Thus, T4 may only need to encode an anti_RM system specifically counteracting type IV R-M immunity. Because most previously characterized ADS and the corresponding database reference models were derived from Caudoviricetes, we repeated the analysis within this class to assess whether the observed pattern was influenced by taxonomic differences in ADS detection. In both the complete viral dataset and the Caudoviricetes subset, viral genome size increased significantly with the number of detected ADS, and viruses encoding ≥2 detectable ADS had substantially larger genomes than those in which no ADS was detected (**Fig. 2b, Fig. S1-S3**). This positive association between viral genome size and ADS copy number further suggests that genome capacity and the evolutionary costs of maintaining multiple immune-evasion modules may constrain viral counter-defense repertoires[64].

Genomic context analysis further revealed that ADS co-occurrence patterns are highly structured in large viral genomes. The most frequent combinations involved different subtypes within anti_RM, anti_CRISPR, and anti_Thoeris families (**Fig. 2c, Fig. S4**), suggesting repeated co-retention of functionally related counter-defense systems. Several larger viral genomes also encoded exceptionally complex ADS repertoires containing multiple distinct ADS classes, including combinations of anti_CBASS, anti_DarTG, anti_RM, and anti_TA systems (**Fig. 2d**). These findings suggested that viruses with large genomes may deploy coordinated multi-layered counter-defense strategies to overcome hosts carrying complex and multilayered immune systems.

### Viral lifestyle determines the deployment of anti-defense strategies across infection stages

ADS identified from the IMG/VR and VIRE datasets targeted diverse prokaryotic antiviral defense pathways. Systems including anti_CRISPR, anti_Dnd, anti_RM, anti_RecBCD, and anti_BREX primarily counteracted defenses targeting invading viral nucleic acids[65], whereas anti_CBASS, anti_Thoeris, NADP, anti_Gabija, anti_Retron, anti_Pycsar, anti_TA, anti_DarTG, and anti_DSR were mainly interfered with abortive infection and immune signaling pathways activated during middle and later stages[27, 66]. This functional partitioning suggests that viruses encounter distinct immune barriers across different stages of infection, raising the possibility that viral lifestyles shape the deployment of corresponding counter-defense strategies.

We next classified ADS-carrying viral genomes according to predicted lifestyle. In IMG/VR, ADS targeting DNA recognition and cleavage systems, including anti_CRISPR, anti_RM, and anti_Dnd, were preferentially associated with temperate viruses, with more than half of these systems occurring in genomes of temperate viruses (**Fig. 3a, Supplementary Text S2**). In contrast, ADS associated with abortive infection and immune signaling pathways, including anti_CBASS, anti_TA, and NADP, were predominantly enriched in virulent viruses (**Fig. 3a**). Similar partitioning patterns were also observed at the community level. Approximately 70% of ADS detected in temperate viruses targeted nucleic acid immunity, whereas virulent viruses allocated a substantially larger fraction of their ADS repertoire to systems associated with abortive infection and immune signaling pathways (**Fig. 3c**). Consistent lifestyle-associated patterns were independently recovered in the VIRE dataset (**Fig. 3b,c**). To reduce the potential confounding effect of host taxonomy, we further examined temperate and virulent viruses infecting Pseudomonadota and Bacillota, the two host phyla with the largest sample sizes. The same lifestyle-associated differences in ADS composition were observed within both host groups (**Fig. S5)**, indicating that the overall pattern was not driven solely by differences in host taxonomic composition. These patterns support distinct immune barriers encountered during alternative infection strategies. Establishing stable lysogeny require temperate viruses to evade sequence-specific host defenses[31], whereas virulent viruses undergoing rapid intracellular replication are likely more vulnerable to middle- and later-stage immune pathways capable of disrupting infection progression[67]. Together, these results indicate that viral infection strategy structure viral ADS repertoires.

**Fig. 3.**
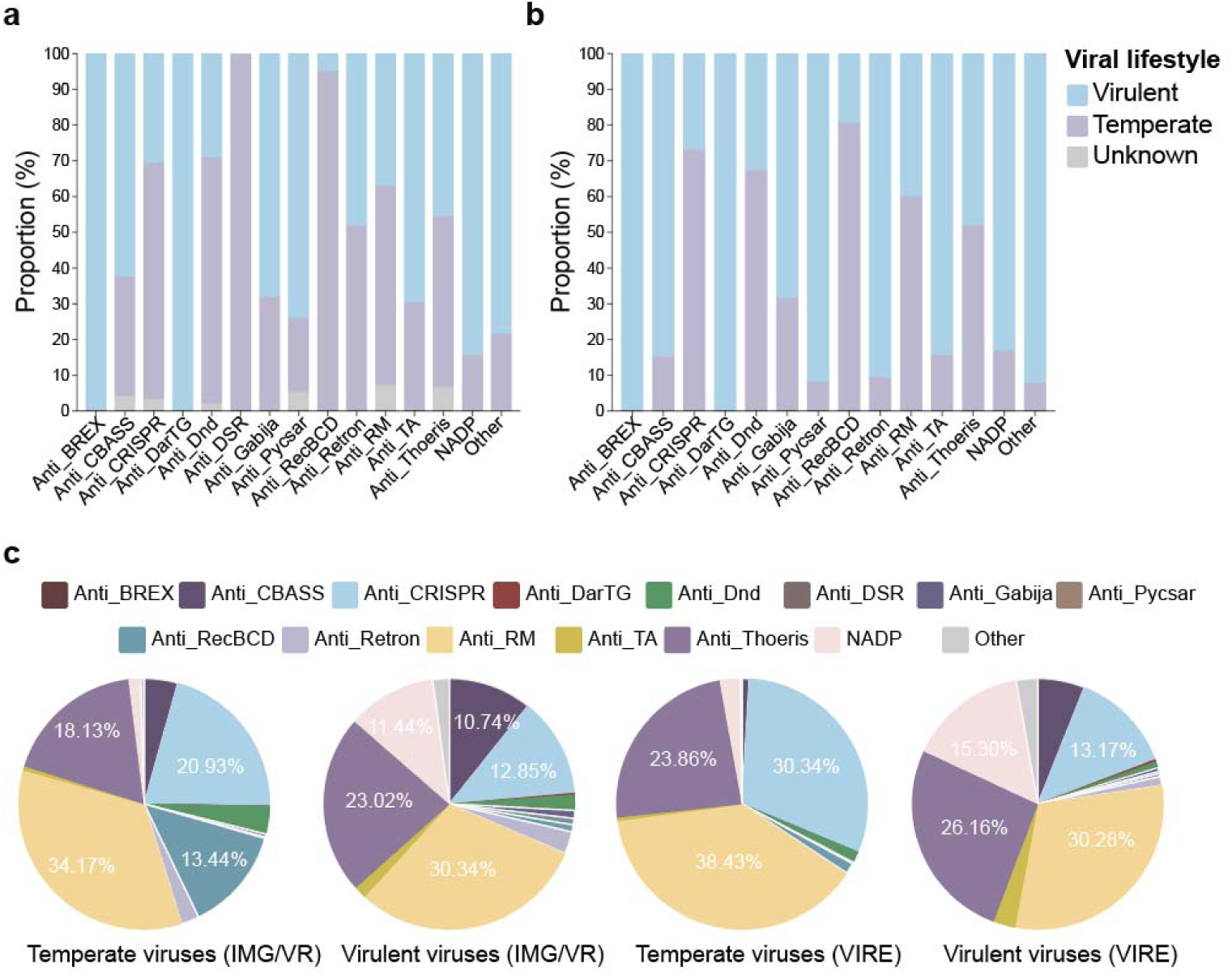
Viral lifestyle shapes the deployment of ADS across infection strategies. Distribution of ADS classes among temperate, virulent, and unclassified viral populations in the IMG/VR database (**a**) and the VIRE database (**b**). (**c**) Relative composition of ADS classes in temperate and virulent viruses from the IMG/VR database and the VIRE database.

### Prokaryotic host-associated structuring of viral anti-defense repertoires

Given that ADS evolve in response to specific host immune barriers [68], ADS repertoires may also be shaped by host defenses. Notably, prokaryotic genomes often encode multiple antiviral defense systems, and their compositions vary substantially across microbial lineages[37, 68]. For example, CRISPR-Cas systems occur in only ∼40% of bacterial genomes[69]. Such heterogeneity in host immunity is therefore expected to impose lineage-specific selective pressures on viral ADS. We therefore analyzed ADS-encoding viral genomes with host annotations provided in the IMG/VR database to test whether viruses infecting different host lineages encode distinct anti-defense repertoires.

A total of 35 host-associated phyla were identified for ADS-carrying viruses, among which Pseudomonadota (44.34%), Bacillota (32.44%), and Bacteroidota (20.11%) were dominant (**Fig**. **4a**). Viral ADS repertoires differed markedly among host-associated viruses (**Fig. 4b, Supplementary Text S3**). Viruses associated with Pseudomonadota exhibited the most diverse ADS repertoires, whereas viruses linked to Bacteroidota were strongly enriched in anti_RM systems. In Bacillota-associated viruses, anti_CRISPR and anti_Thoeris predominated, whereas Fusobacteriota- and Thermoproteota-associated viruses showed extreme enrichment of anti_CRISPR systems. Similar lineage-associated partitioning was also observed at the genus level (**Fig. S6**). For example, *Escherichia*-associated viruses were enriched in anti_RecBCD and anti_RM systems, whereas anti_CRISPR overwhelmingly dominated viruses linked to *Staphylococcus* and *Streptococcus*. These lineage-associated patterns are consistent with previous observations that antiviral defense systems are unevenly distributed across prokaryotic taxa. For example, restriction-modification systems are widespread and highly diversified in Bacteroidota[70], whereas CRISPR-Cas systems are prevalent in many *Streptococcus* lineages[71]. These results indicated that viral ADS repertoires are strongly shaped by host phylogeny and their immune architecture. The uneven distribution of prokaryotic defense systems across microbial taxa likely creates distinct immune-selection landscapes that favor the maintenance of different viral ADS in different host lineages. Consequently, environmental changes that alter microbial community composition and host immune landscapes may further drive the shifts in viral ADS repertoires across ecosystems.

**Fig. 4.**
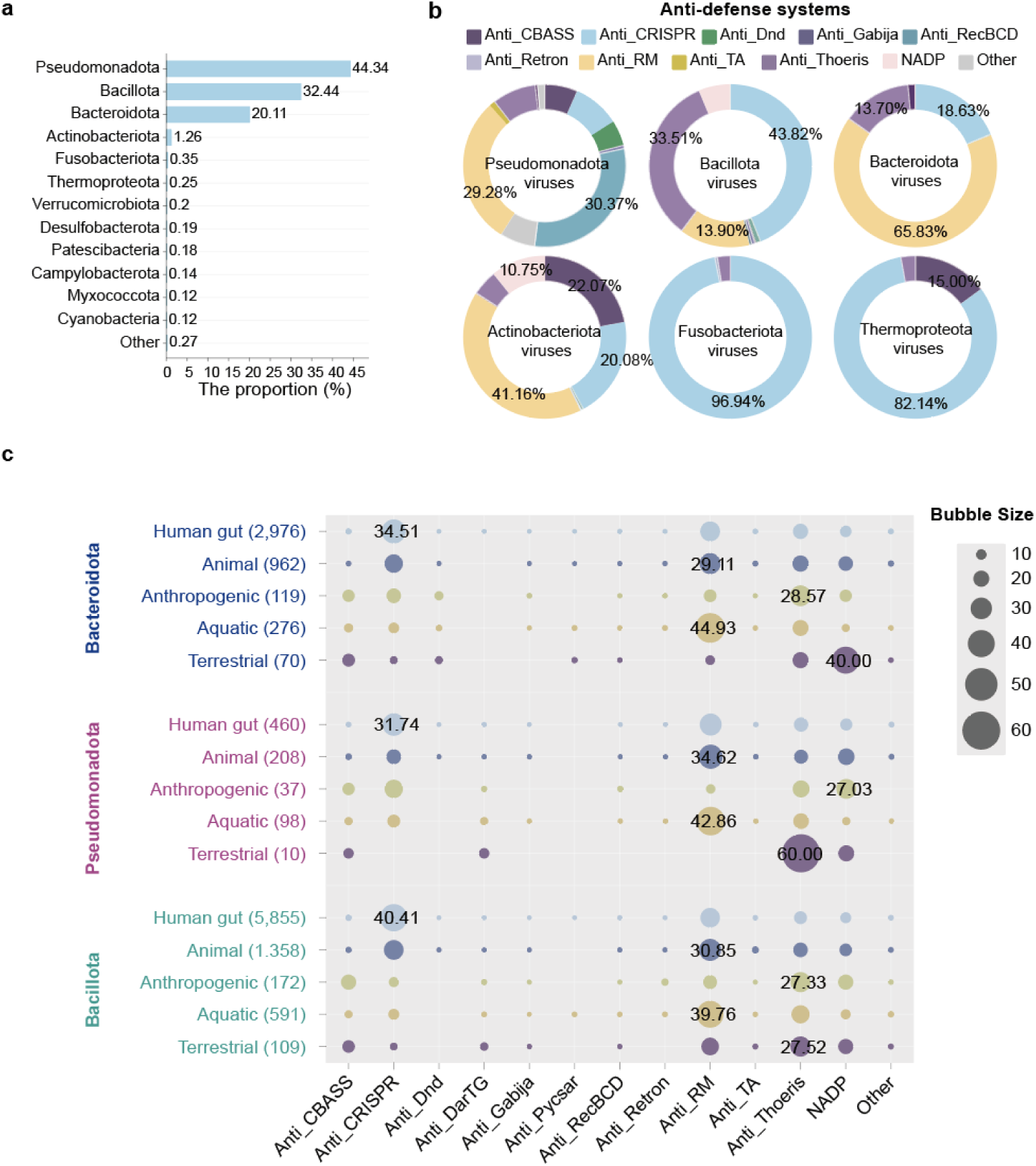
Host-associated structuring of viral ADS repertoires. (**a)** Distribution of predicted host phyla for ADS-carrying viruses in the IMG/VR database. Only phyla accounting for more than 0.01% of total host assignments are shown individually, whereas the remaining low-abundance phyla are grouped as “Other”. (**b**) Composition of ADS repertoires in viruses associated with the six most prevalent host phyla, including Pseudomonadota, Bacillota, Bacteroidota, Actinobacteriota, Fusobacteriota, and Thermoproteota. Pie charts show the relative proportions of different ADS classes within each host-associated viral group. (**c**) Habitat-associated composition of detectable ADS classes in VIRE viruses infecting hosts from Bacteroidota, Pseudomonadota, and Bacillota. Rows represent viral groups from the human gut, other animal-associated environments, anthropogenic environments, aquatic environments, and terrestrial environments. Numbers in parentheses indicate the number of ADS-carrying viral genomes in each group. Bubble size represents the relative proportion of each ADS class within the corresponding host phylum–habitat group; selected proportions are labeled. Only viral genomes with >90% estimated completeness were included in the analysis.

### Habitat-associated structuring of viral anti-defense repertoires

Distinct habitats harbor divergent prokaryotic community composition and viral lifestyles, which are likely to drive systematic differentiation of viral ADS repertoires. To test this, we compared ADS repertoires across human gut-associated, animal-associted, anthropogenic, aquatic, and terrestrial environments using viral genomes from the VIRE database. The prevalence of ADS varied markedly among habitats (**Fig. S7**). Viral genomes carrying at least one ADS, particularly those encoding multiple ADS, were more frequent in anthropogenic and terrestrial environments than in aquatic or human gut-associated systems. The accumulation and combinatorial deployment of viral ADS, potentially reflect intensified virus-host coevolution in terrestrial and anthropogenically impacted ecosystems[3, 30, 72, 73].

Across habitats, viral ADS repertoires retained a conserved core of dominant ADS classes, although their relative composition differed substantially among environments (**Fig. S7, Supplementary Text S4**). Terrestrial and anthropogenic environments were both enriched in anti_Thoeris-dominated repertoires (32.53% and 28.85%), whereas human gut-associated, animal and aquatic viral communities showed comparatively stronger enrichment of anti_RM systems (33.90%, 36.94% and43.51%). Human gut-associated viral communities were additionally enriched in anti_CRISPR systems (25.87%), consistent with the high prevalence of lysogeny and CRISPR-Cas immunity in host-associated microbiomes. In contrast, anthropogenic environments exhibited more balanced ADS repertoires, potentially reflecting the ecological mixing of host-associated and natural microbial communities[74]. Because viral ADS repertoires may also vary with host taxonomy, we further examined habitat-associated patterns separately for viruses infecting Bacteroidota, Pseudomonadota, and Bacillota. Habitat-associated shifts in ADS composition remained evident within each host phylum (**Fig. 4c**), indicating that these patterns could not be explained solely by differences in host-community composition among habitats. Collectively, these findings indicated that habitat acts as an important ecological filter shaping viral ADS strategies. Variation in host community composition, immune architectures, and virus-host interaction dynamics[28] across ecosystems likely contributes to the systematic differentiation of viral ADS repertoires.

### Heavy metal stress is associated with shifts in virus-host immune interactions

Building on the habitat-associated shaping of ADS repertoires, we investigated whether viral ADS and host defense profiles varied along environmental stress gradients.. Heavy metal contamination imposes strong oxidative and physiological stress on microbial communities and has been associated with elevated viral abundance and provirus induction[50, 75]. We therefore examined whether such abiotic pressure could alter ADS and their interactions with host immunity. To address this question, we analyzed 401 soil metagenomes spanning a gradient of heavy metal contamination across China. Pollution intensity was quantified using the Nemerow pollution index (P ) based on seven metals (Cd, As, Pb, Cr, Cu, Ni, and Zn; see **Methods**). Samples were classified as clean soils (n = 92), slightly contaminated soils (n = 139), or heavily contaminated soils (n = 170), representing increasing levels of metal contamination (**Fig. 5a**).

**Fig. 5.**
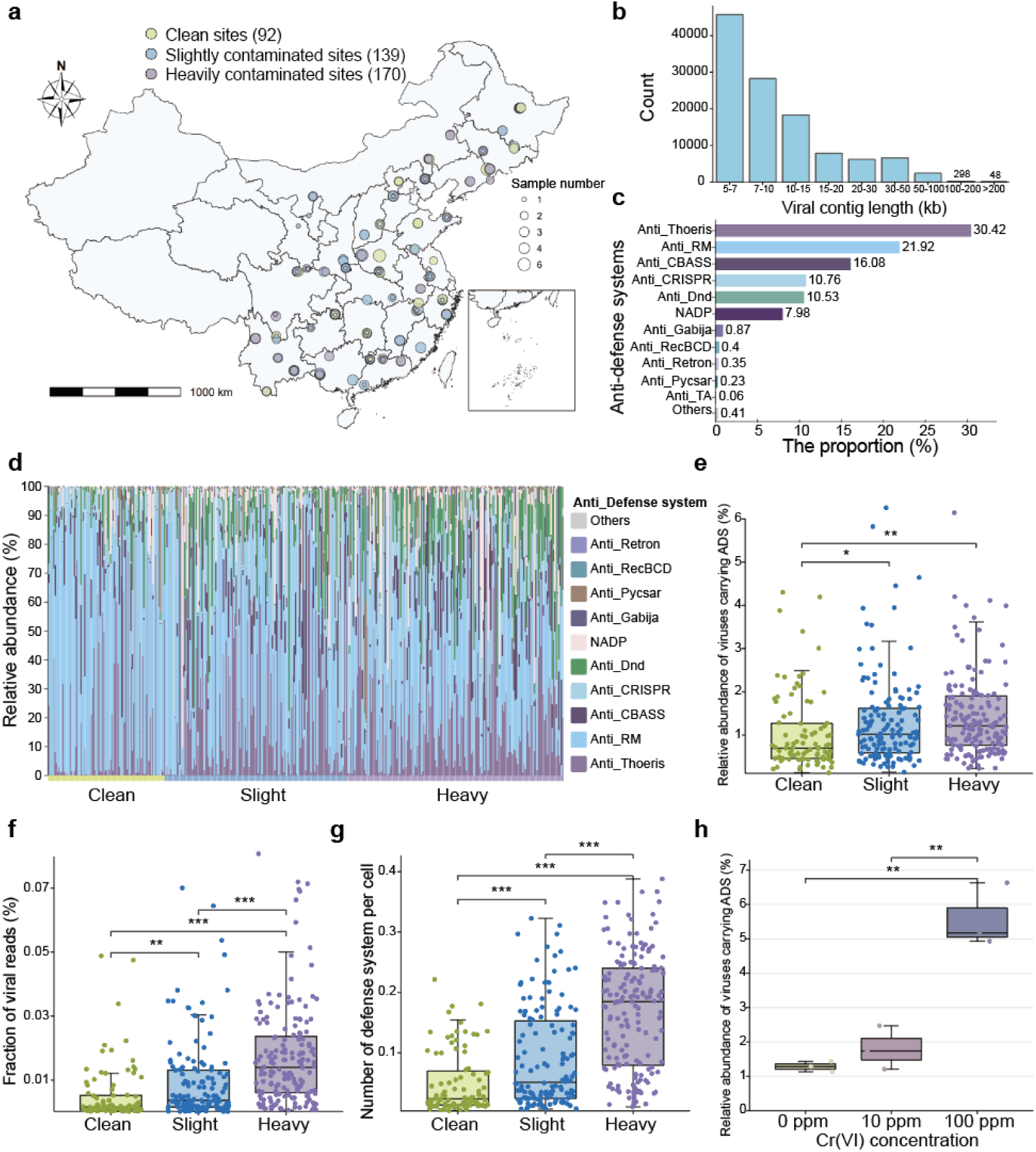
Environmental stress-associated structuring of viral ADS repertoires. **(a)** Soil metagenomes were collected along a gradient of heavy metal contamination. Pollution intensity was quantified using the Nemerow pollution index (P ) based on seven metals (Cd, As, Pb, Cr, Cu, Ni, and Zn). Samples were classified into three categories: clean soils (P ≤ 0.7, n = 92; green dots), slightly contaminated soils (0.7 < P ≤ 10.0, n = 139; blue dots), and heavily contaminated soils (P > 10.0, n = 170; purple dots), representing increasing levels of metal stress. **(b)** Length distribution of viral contigs. **(c)** Composition of ADS encoded by viral communities in heavy metal contaminated soil. **(d)** Shifts in the composition of ADS under increasing heavy metal stress. **(e)** Changes in the relative abundance of ADS-carrying viruses along the heavy metal contamination gradient. **(f)** Changes in the relative abundance of viral reads under heavy metal contamination. **(g)** Variation in the number of ADS per prokaryotic genome across the contamination gradient. **(h)** Changes in the relative abundance of ADS-carrying viruses with increasing chromium concentrations in microcosm experiments. (Wilcoxon rank-sum test, ***, *P* < 0.001, **, *P* < 0.01, *, *P* < 0.05)

We recovered 115,572 viral contigs (>5 kb) from heavy metal contaminated soil (see **Fig. 5b, Supplementary Text S5**), including 48 jumbo phage genomes (>200 kb), and identified 1,729 ADS spanning 13 major classes and 40 subtypes (**Fig. 5c, Fig. S8-S10**). Lifestyle-associated ADS partitioning patterns and genome size constraints were also conserved under metal stress (**Fig. S11-S13**). As in global terrestrial viral communities, anti-Thoeris (30.42%) and anti-RM (21.92%) were the predominant ADS types. Notably, among ADS-carrying viruses, increasing heavy metal stress was accompanied by a compositional shift from anti_RM-dominated repertoires toward a more even distribution across multiple ADS classes, with the proportion of anti_Thoeris-carrying viruses showing the most pronounced increase (**Fig. 5d**).

The proportion of ADS-carrying viruses increased from 1.0% in clean soils to 1.7% in heavily contaminated soils (*P* < 0.01, **Fig. 5e**). Viral relative abundance increased approximately fourfold from clean to heavily contaminated soils (*P* < 0.01, **Fig. 5f**), accompanied by significant increases in viral richness and diversity (*P* < 0.01, **Fig. S14**), indicating enhanced viral proliferation under metal stress. In parallel, prokaryotic antiviral defense levels were also strongly enriched, with both defense system abundance and defense-related genes increasing significantly in contaminated soils (*P* < 0.01, **Fig. 5g, Fig. S15**). Moreover, Prokaryotic community composition also differed across contamination levels. For example, from clean to heavily contaminated soils, the relative abundance of Actinomycetota decreased at the phylum level, whereas that of *Sphingomonas* increased at the genus level (**Figs. S15** and **S16**). Thus, the higher proportion of ADS-carrying viruses along the contamination gradient may reflect concurrent shifts in prokaryotic community composition and their antiviral defense-system profiles, alongside an overall increase in viral abundance.

To independently validate whether these field-associatedpatterns extend to other heavy metals, we further conducted chromium-stress soil microcosm experiments under controlled conditions, with three contamination levels (Cr(VI) = 0, 10, and 100 mg/kg (ppm); see Methods). Because fragmented viral assemblies may substantially limit the recovery of viral ADS in environmental viromes, we applied a hybrid assembly strategy combining short- and long-read sequencing. Compared with short-read assembly alone, hybrid assembly substantially improved viral genome recovery by 24.7% and ADS detection sensitivity by 51.7% (see **Supplementary Text S6**). The relative abundance of ADS-carrying viruses increased significantly with Cr(VI) concentration (*P* < 0.01, **Fig. 5h**), accompanied by a parallel increase in prokaryotic antiviral defense systems[50], thereby recapitulating the coordinated patterns observed in the field dataset. Together, these concordant changes are consistent with shifts in virus–host immune dynamics under metal stress and suggest that metal-stressed ecosystems may provide valuable settings for investigating microbial defense and counter-defense systems.

### ADS enrichment may contribute to microbial adaptive potential under metal stress

ADS can suppress prokaryotic immune barriers that also restrict horizontal gene transfer[21, 24]. We therefore examined whether the occurrence of viral ADS was associated with HRG and other mobile genetic elements. HRGs were detected more frequently in ADS-carrying viruses than in viruses without detected ADS (1.85% vs 0.43%; **Fig. 6a,b**). This association persisted after accounting for viral contig length (adjusted OR = 1.86, *P* = 0.003). This co-occurrence suggests an association between ADS carriage and the genetic potential for heavy metal resistance.

**Fig. 6.**
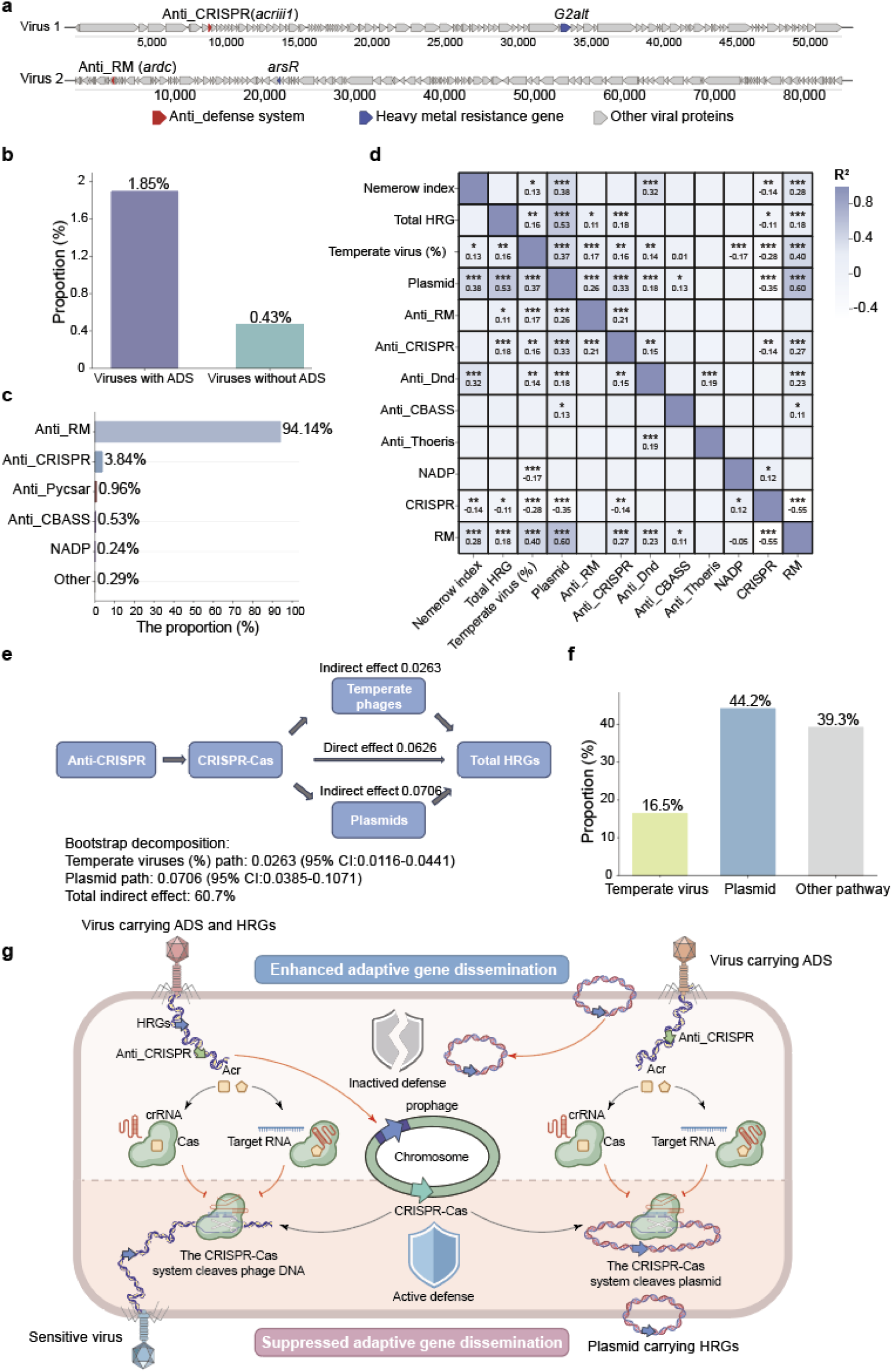
Ecological implications of ADS enrichment on microbial adaptive capacity under metal stress. (**a**) Representative genome architectures of viruses co-encoding ADS and heavy metal resistance genes (HRGs). (**b**) Proportion of virsl genomes carrying HRGs in ADS-positive and ADS-negative groups, showing a higher frequency of HRG occurrence in ADS-containing viruses. (**c**) Distribution of ADS types encoded by plasmids, with anti_RM systems dominating the ADS repertoire. (**d**) Correlation analysis among ADS, host defense systems, and HRG abundance based on Pearson correlations; color intensity represents correlation strength (R²), with asterisks indicating statistical significance (***, *P* < 0.001, **, *P* < 0.01,*, *P* < 0.05). (**e**) Parallel mediation framework illustrating the relationship between anti_CRISPR and total HRG abundance, with temperate virus proportion and plasmid abundance modeled as parallel mediators. (**f**) Decomposition of the total effect of anti_CRISPR on HRG abundance based on causal mediation analysis, showing the relative contributions of the temperate virus-mediated pathway, plasmid-mediated pathway, and the remaining other effects. (**g**) Potential mechanisms by which ADS-carrying temperate viruses promote the dissemination and enrichment of adaptive genes in microbial communities. Viruses that carry both ADS and HRGs may overcome host defense systems and persist as prophages, thereby enriching HRGs. In addition, ADS-carrying viruses may suppress host defense systems, facilitating the dissemination and stable maintenance of HRG-carrying plasmids.

On the other hand, similar anti-defense profiles dominated by anti_RM and anti_CRISPR systems were also observed in plasmids (**Fig.6c**), suggesting that immune suppression-associated ADS may be shared across multiple mobile genetic elements. At the community level, the relative abundance of anti_CRISPR-carrying viruses was positively associated with both plasmid abundance and the proportion of temperate viruses (*P* < 0.001, **Fig. 6d**). These relationships are consistent with coordinated variation among viral counter-defense functions and mobile genetic elements in metal-stressed microbial communities. We further used mediation analysis to explore statistical pathways linking anti-CRISPR abundance with total HRG abundance. Anti_CRISPR abundance was positively associated with plasmid abundance (β = 0.2784, *P* = 1.86 × 10 ) and the proportion of temperate viruses (β = 0.0284, *P* < 0.001). In the fitted mediation model, the statistical pathway through plasmid abundance accounted for 44.2% of the estimated total association, whereas the pathway through the proportion of temperate viruses accounted for 16.5% (**Fig. 6e-g**). Together, these findings raise the possibility that, beyond their roles in viral infection, ADS may form part of a broader counter-immunity network associated with mobile genetic element dynamics and the distribution of adaptive traits in stressed microbial communities. However, whether ADS directly influence these processes remains to be established experimentally.

## Conclusion

Our study reveals consistent community-level patterns in viral ADS repertoires across diverse ecosystems. ADS repertoire size and composition varied systematically with viral genome size, lifestyle, host lineage, and habitat, linking viral anti-defense strategies to their genomic and ecological context. These patterns extended to metal-contaminated soils, where changes in ADS prevalence and composition accompanied shifts in prokaryotic defense profiles and community composition. Chromium-stress microcosms further supported the association between metal exposure and increased abundance of ADS-carrying viruses. Associations of viral ADS with mobile genetic elements and heavy metal resistance genes also suggest ecological connections between anti-defense and the distribution of adaptive traits.

By extending anti-defense research beyond individual virus–host systems, these findings bring viral anti-defense into the study of microbial community ecology. Considering ADS alongside host defenses offers a more integrated perspective on virus–host immune interactions, complementing descriptions based on community composition and viral abundance. This perspective provides a basis for investigating how immune interactions relate to viral persistence, genetic exchange, and microbial responses to environmental disturbance. It also identifies questions relevant to microbiome management, including how anti-defense characteristics can inform virus selection and the assessment of mobile genetic element dissemination.

The ecological patterns reported here are based on currently detectable ADS families and may therefore be affected by uneven family coverage, despite our efforts to minimize this bias[37]. Expanding reference databases and developing more broadly applicable detection approaches will help capture the diversity of viral anti-defense systems more comprehensively. Future studies incorporating direct measurements of virus–host interactions and horizontal gene transfer will be essential for testing the functional implications of our findings and linking community-level ecological patterns to the molecular mechanisms underlying microbial immune ecology.

## Author contributions

Conceptualization: D.H., P.F.Y. conceived the project and designed experiments. D.H. performed experiments, analyzed data, and wrote the manuscript. P.F.Y. acquired funding and revised the manuscript. Y.Y.Y. participated in the analysis and discussion of the results. All authors edited the manuscript.

## Consent for publication

All the authors approved the submission.

## Data availability

The two global viral genome resources used in this study were the IMG/VR[34] and VIRE[35] datasets. Metagenomic sequencing data from heavy metal-contaminated soils in China are available from the NCBI BioProject database under accession numbers PRJNA1253350, PRJNA1253357, PRJNA1253358, PRJNA1253360, and PRJNA1254387. Metagenomic sequencing data from the microcosm experiments are available from the NCBI BioProject database under accession number PRJNA1104006.

