## Supplemental Figures and Texts for "Anti-defense systems of viral communities vary with genomic traits, host immunity and environmental stress"

**This study includes 6 supplementary texts and 17 supplementary figures.**

**Supplementary text S1: The composition of viral ADS repertoires in IMG/VR and VIRE datasets.**

A total of 101,594 ADS were identified in 92,211 IMG/VR viral genomes (1.64%) and 59,801 ADS were identified in 54,465 VIRE viral genomes (6.80%), spanning 14 and 13 major ADS classes, respectively. Viral ADS repertoires exhibited remarkably consistent composition patterns. In both IMG/VR and VIRE, anti_RM (33.78% versus 34.76%), anti_Thoeris (20.89% versus 24.89%) and anti_CRISPR (17.17% versus 22.48%) represented the dominant ADS categories, together accounting for more than 70% of all detected systems. (Fig. 1A,1B). In IMG/VR, the next most abundant ADS classes included anti_RecBCD (7.48%), anti_CBASS (6.69%), NADP (5.39%), and anti_Dnd (2.79%). By contrast, VIRE viral genomes were additionally enriched in NADP (8.42%), anti_CBASS (3.26%), anti_TA (1.55%), and anti_Dnd (1.13%). The convergence of ADS composition across independent global viral datasets suggests that viral ADS repertoires may be shaped by common ecological and evolutionary constraints across viral communities. Moreover, in the IMG/VR dataset, 5,529,187 viral genomes (98.36%) encoded no ADS, 84,898 (1.51%) encoded one ADS, and only 7,313 (0.13%) encoded ≥2 ADS. A similar pattern was observed in VIRE, where 746,675 viral genomes (93.20%) lacked ADS, 50,495 (6.30%) encoded one ADS, and 3,970 (0.50%) encoded ≥2 ADS (Fig. 1C, Fig. S1). Viral genomes carrying different numbers of ADS also showed marked differences in genome size. Viral genome size increased markedly with ADS copy number. In IMG/VR, genomes lacking ADS, carrying one ADS, or encoding ≥2 ADS had average sizes of 14.17 kb, 40.98 kb, and 72.74 kb, respectively, whereas the corresponding values in VIRE were 43.67 kb, 68.32 kb, and 104.96 kb (Fig. 1C, Fig. S1). These results support the idea that genome size is a major constraint on viral anti-defense capacity.

**Supplementary text S2: Temperate and virulent viruses encode different ADS.**

Prevalence of each ADS type in temperate vs. virulent viruses: In IMG/VR, 52.83% of ADS-carrying viral genomes were annotated as temperate viruses, 39.67% as virulent viruses, and 7.50% remained unclassified. ADS targeting DNA recognition and cleavage systems were more frequently associated with temperate viruses than with virulent viruses. For example, anti_CRISPR, anti_RM, and anti_Dnd were predominantly found in temperate viruses, with 66.14%, 55.64%, and 68.87% of these ADS occurring in temperate genomes, respectively (Fig. 2A). In contrast, ADS associated with abortive infection and immune signaling were enriched in virulent viruses. anti_CBASS, anti_TA, and NADP occurred mainly in virulent genomes, with 62.25%, 69.37%, and 84.34% found in virulent viruses, respectively (Fig. 2A).

Relative abundance of ADS categories within temperate vs. virulent viral communities: As expected, in temperate viruses, approximately 70% of detected ADS targeted nucleic acid immunity, including anti_RM (34.17%), anti_CRISPR (20.93%), anti_RecBCD (13.43%), and anti_Dnd (3.58%). By contrast, virulent viruses allocated more than half of their ADS repertoire to abortive infection and immune signaling pathways, including anti_Thoeris (23.02%), NADP (11.44%), and anti_CBASS (10.74%) (Fig. 2C). Among ADS-carrying viral genomes from VIRE database, 55.60% were classified as temperate viruses, 44.04% as virulent viruses, and only 0.16% remained unclassified. Consistent with the IMG/VR results, temperate viruses were preferentially enriched in ADS targeting nucleic acid immunity, whereas virulent viruses more frequently encoded ADS associated with abortive infection and immune signaling pathways (Fig. 2B,2D).

**Supplementary text S3: The composition of ADS is shaped by host immunity.**

As shown in Fig. 3B, ADS repertoires varied markedly among host-associated viruses. At the phylum level (Fig. 3B), viruses associated with Proteobacteria exhibited the most diverse ADS repertoire, encompassing at least 12 ADS classes. anti_RecBCD (30.37%) and anti_RM (29.28%) were the dominant systems, followed by anti_CRISPR (9.36%), anti_Thoeris (8.68%), and anti_Retron (7.06%). In contrast, ADS repertoires in viruses infecting Actinobacteriota were strongly dominated by anti_RM (41.16%), followed by anti_CBASS (22.07%) and anti_CRISPR (20.08%). Distinct host-associated patterns were also observed in other major phyla. Viruses linked to Bacteroidota were overwhelmingly enriched in anti_RM systems (65.83%), with anti_CRISPR (18.63%) and anti_Thoeris (13.70%) representing secondary components. In Firmicutes-associated viruses, anti_CRISPR (43.82%) and anti_Thoeris (33.51%) predominated, whereas anti_RM accounted for a comparatively smaller fraction (13.90%). Fusobacteriota-associated viruses showed an extreme bias toward anti_CRISPR, which represented 96.94% of all detected ADS. Similarly, Thermoproteota-associated viruses were also strongly enriched in anti_CRISPR systems (82.14%), together with a smaller proportion of anti_CBASS (15.00%).

This host-associated variation in ADS repertoires was also evident at the genus level (Fig. 3C). Compared with phylum-level patterns, ADS repertoires within individual host genera were generally less diverse and were often dominated by one or two major ADS classes. Distinct lineage-specific ADS profiles were observed among the most frequently represented host genera. Viruses predicted to infect *Escherichia* were primarily enriched in anti_RecBCD and anti_RM systems, accounting for 43.90% and 36.66% of all ADS detected in *Escherichia*-associated viruses, respectively, with anti_Retron representing an additional 15.21%. In contrast, *Acinetobacter*-associated viruses mainly encoded anti_Thoeris (49.69%), anti_Dnd (25.10%), and anti_Retron (16.94%). Among Firmicutes-associated genera, anti_CRISPR overwhelmingly predominated in viruses linked to *Staphylococcus* and *Streptococcus*, accounting for 95.03% and 98.08% of detected ADS, respectively. Collectively, these results indicate that ADS composition is strongly shaped by host phylogeny, and demonstrate that host immune architecture shapes viral ADS repertoires.

**Supplementary text S4: The composition of ADS is shaped by habitat.**

To test the influence of habitat on ADS distribution, we compared ADS repertoires across five habitats from the VIRE database, including viral genomes from terrestrial environments (51,765), aquatic systems (126,633), the human gut (657,315), and anthropogenic environments (23,358). The prevalence of ADS varied markedly across habitats (Fig. 3D). Viral genomes carrying at least one ADS were more abundant in anthropogenic (6.60%) and terrestrial habitats (5.60%) than in human gut (5.30%) and aquatic systems (4.60%). Similarly, among ADS-carrying viral genomes, genomes encoding multiple ADS were most frequent in anthropogenic environments (14.40%), followed by terrestrial (13.50%), aquatic (8.20%), and human gut environments (6.70%). Among, terrestrial environments were primarily characterized by anti_Thoeris (29.70%), followed by anti_RM (17.50%), whereas aquatic systems were strongly dominated by anti_RM (40.30%) together with a secondary enrichment of anti_Thoeris (23.20%). Human gut-associated viral communities showed co-enrichment of anti_RM (33.90%) and anti_CRISPR (25.90%), with anti_Thoeris representing an additional 24.90%. Anthropogenic environments exhibited a more balanced distribution of anti-defense systems, with anti_Thoeris (28.90%) as the dominant system, followed by anti_RM (17.20%), anti_CBASS (17.00%), and anti_CRISPR (15.00%).

**Supplementary text S5: The profiles of viral community in heavy metal contaminated soils.**

Taxonomic annotation assigned 61,974 sequences to known viral taxonomy, whereas more than 46% remained unclassified, indicating substantial unexplored viral diversity in soils. Among classified viruses, members of *Caudoviricetes* dominated across nearly all samples, comprising approximately 30% of the viral community (Fig. S4).

**Supplementary text S6: hybrid assembly of short- and long-read improve recovery of viral genomes and ADS.**

To address that soil viral genomes are more likely to be incomplete, we managed to improve recovery of viral genomes and associated defense modules by applying a hybrid sequencing approach combining short- and long-read data in microcosm. Compared with short-read assembly alone, hybrid assembly increased the number of recovered viral contigs (>5 kb) by 24.65% (from 4,309 to 5,371) and the number of detected ADS by 51.69% (from 89 to 134), with the ADS carriage rate rising from 2.07% to 2.50%, demonstrating substantially improved detection sensitivity.

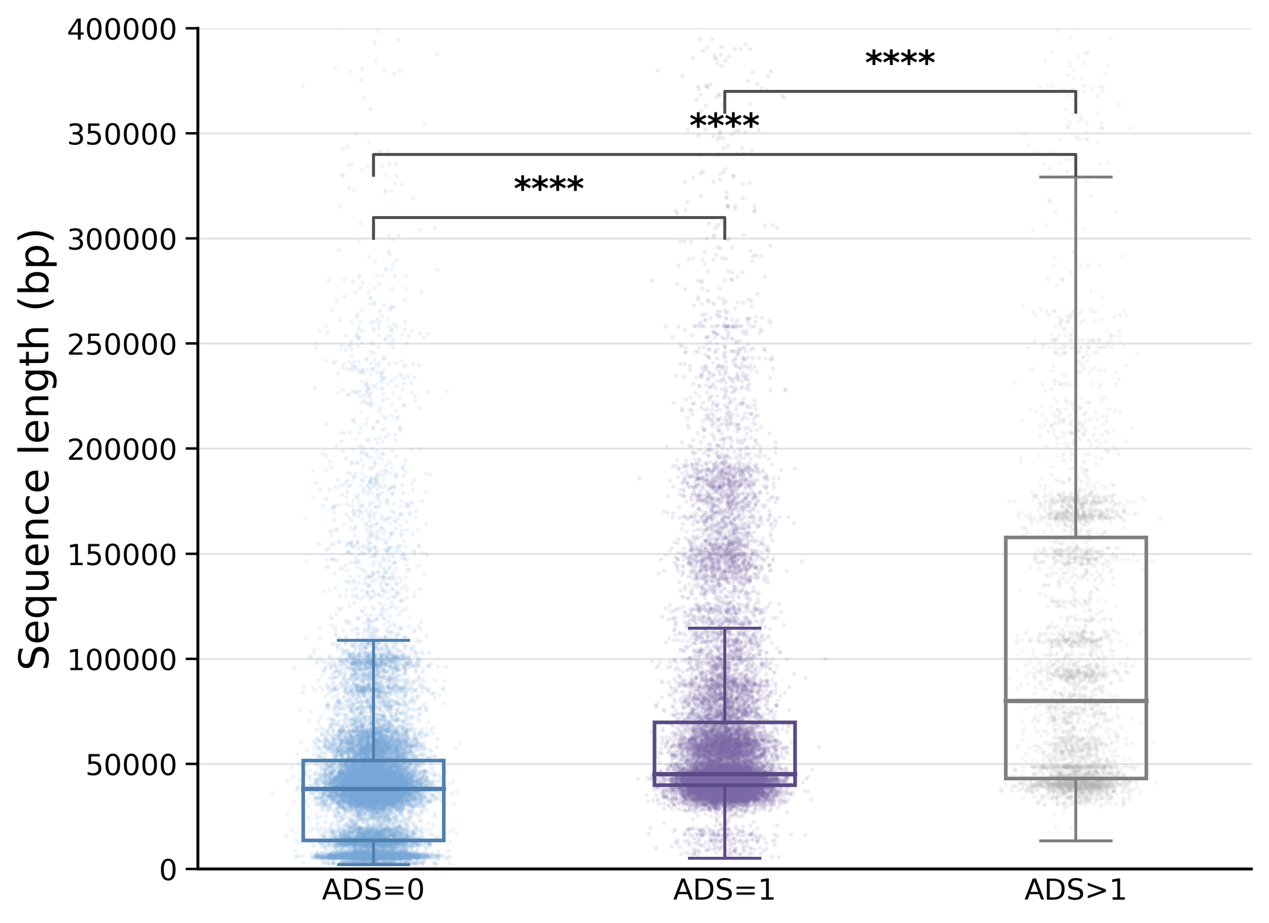

**Figure S1. Genome size distributions of viral genomes carrying different numbers of ADS in VIRE database.** Viral genomes were grouped according to the number of encoded ADS (0, 1 or ≥2 ADS). Boxplots show the distribution of viral genome lengths within each category. Center lines indicate medians, boxes indicate interquartile ranges (IQRs), and whiskers represent 1.5 × IQR. Statistical significance was assessed using two-sided Wilcoxon rank-sum tests (****, *P* < 0.0001).

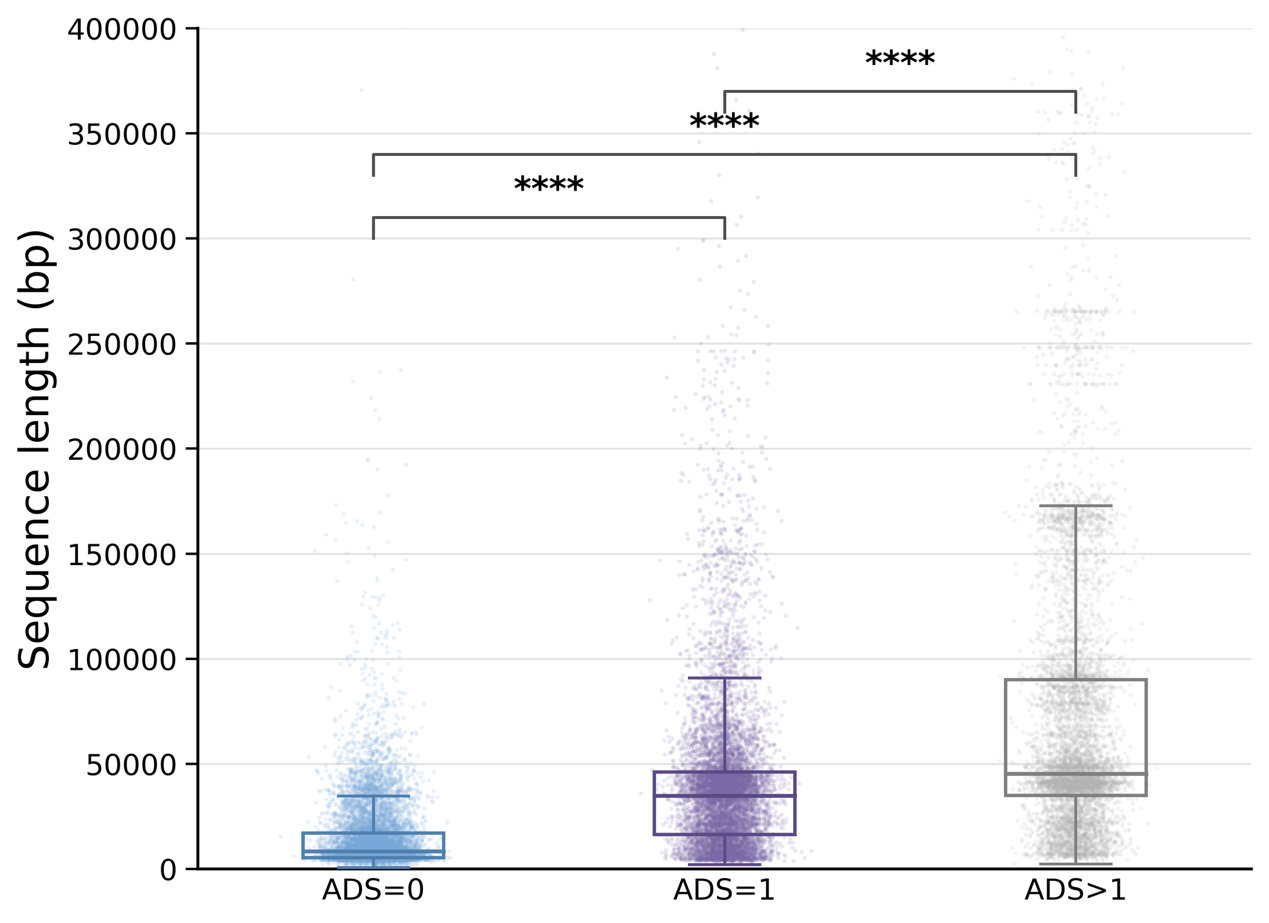

**Figure S2. Genome size distributions of Caudoviricetes genomes carrying different numbers of ADS in IMG/VR database.** Viral genomes were grouped according to the number of encoded ADS (0, 1 or ≥2 ADS). Boxplots show the distribution of viral genome lengths within each category. Center lines indicate medians, boxes indicate interquartile ranges (IQRs), and whiskers represent 1.5 × IQR. Statistical significance was assessed using two-sided Wilcoxon rank-sum tests (****, *P* < 0.0001).

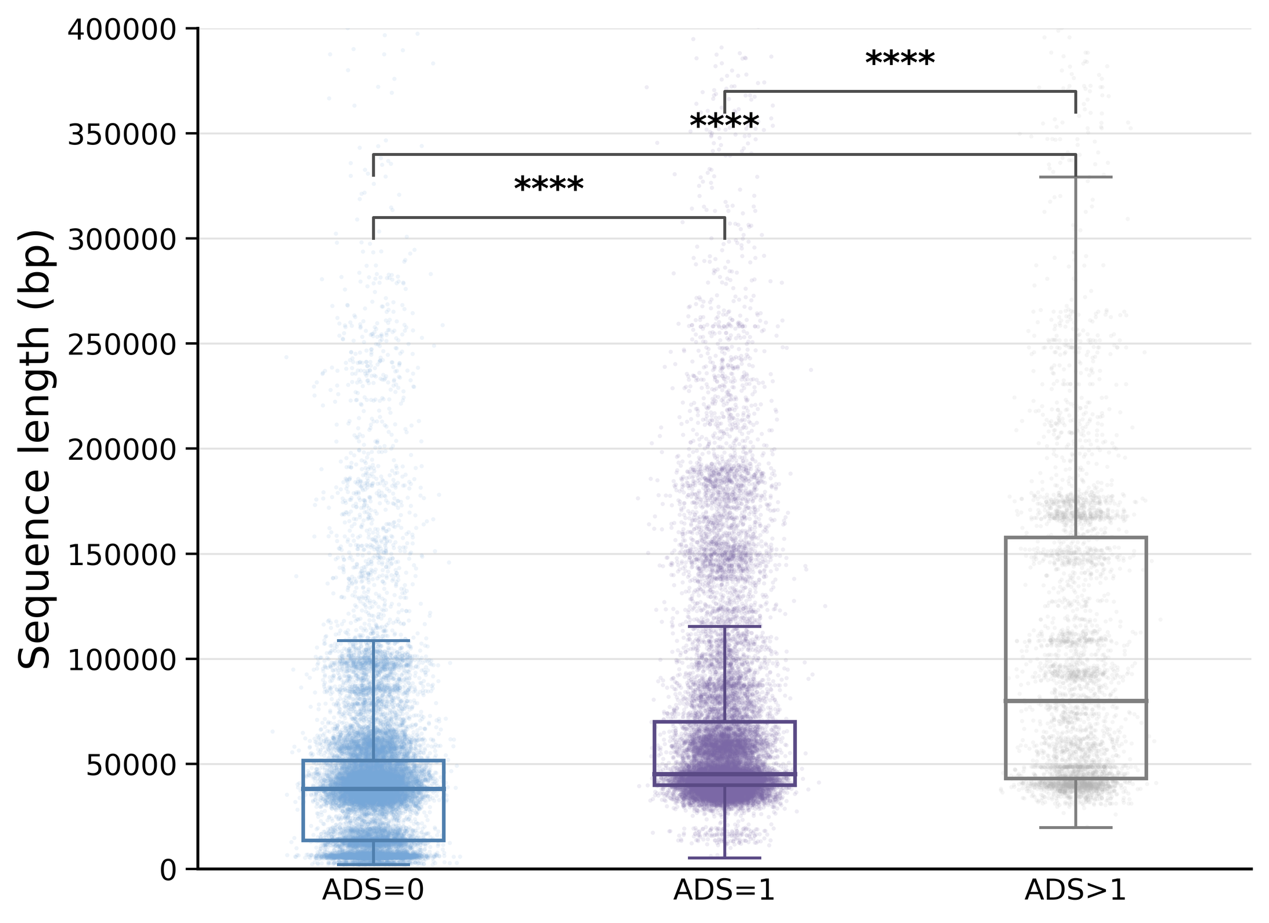

**Figure S3. Genome size distributions of Caudoviricetes genomes carrying different numbers of ADS in VIRE database.** Viral genomes were grouped according to the number of encoded ADS (0, 1 or ≥2 ADS). Boxplots show the distribution of viral genome lengths within each category. Center lines indicate medians, boxes indicate interquartile ranges (IQRs), and whiskers represent 1.5 × IQR. Statistical significance was assessed using two-sided Wilcoxon rank-sum tests (****, *P* < 0.0001).

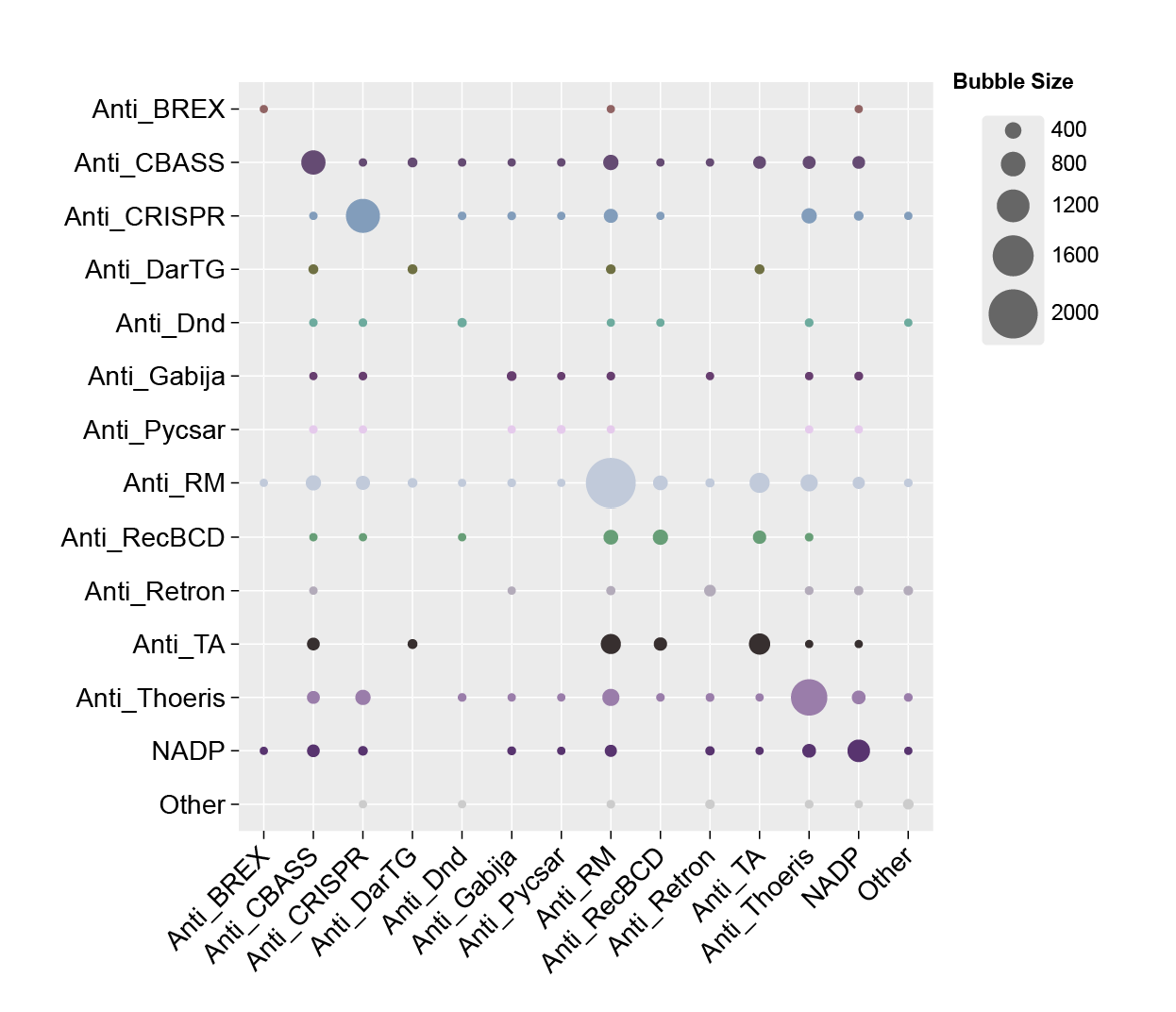

**Figure S4. Pairwise co-occurrence frequencies of ADS within the same viral genomes in the VIRE database.** Bubble sizes indicate the number of viral genomes carrying each ADS combination, highlighting preferential co-occurrence patterns among major ADS classes.

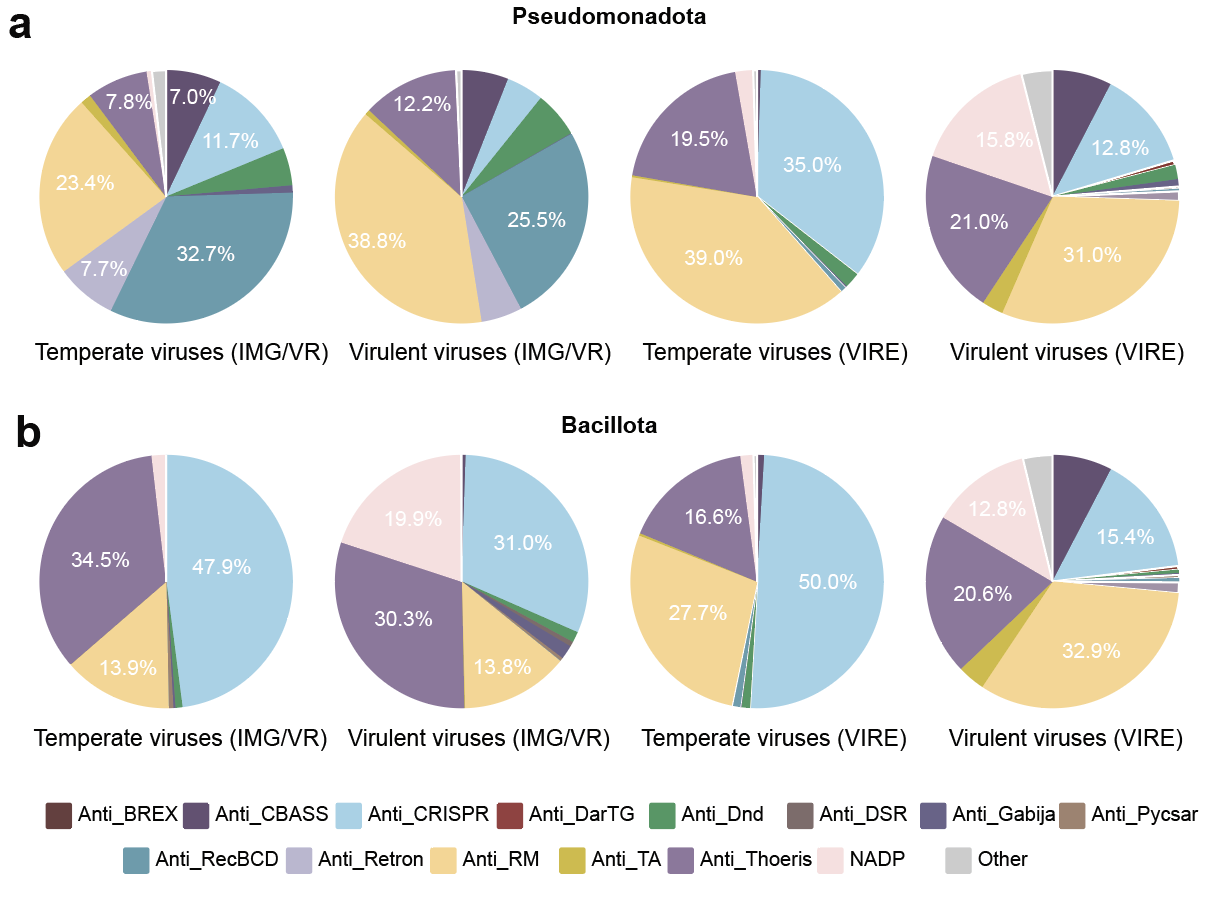

**Figure S5. Relative composition of ADS classes in temperate and virulent viruses infecting Pseudomonadota (a) and Bacillota (b), based on viruses from the IMG/VR and VIRE databases.**

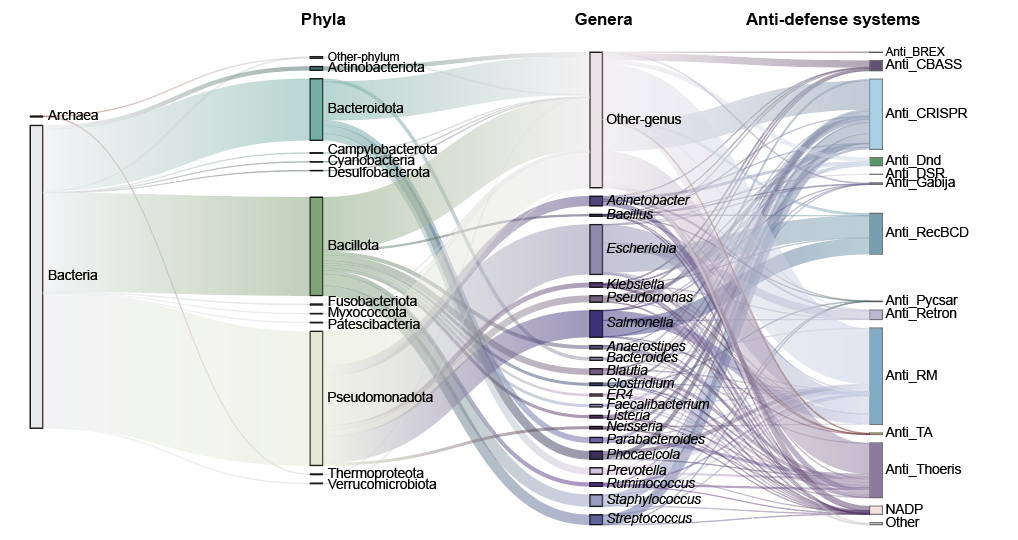

**Figure S6. Sankey diagram illustrating the hierarchical relationships among host domains, phyla, genera, and ADS classes in ADS-carrying viruses.** Flow widths are proportional to the number of associated viral genomes. Only the most abundant host genera and ADS classes are displayed individually, whereas low-abundance categories are grouped as “Other-genus” or “Other”.

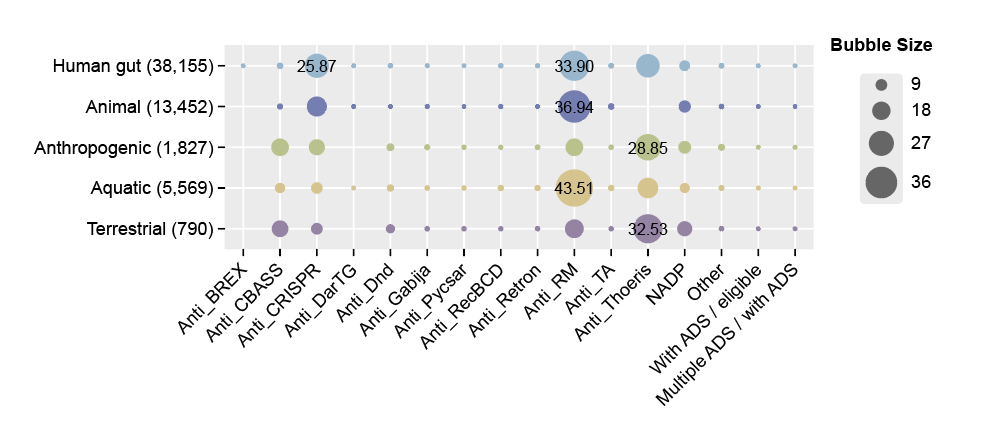

**Figure S7. Prevalence and composition of ADS in viral communities across different habitats from VIRE database.** ADS repertoires across five habitats from the VIRE database, including viral genomes from human gut (657,315), animal (196,143), anthropogenic environments (23,358). aquatic systems (126,633), and terrestrial environments (51,765).

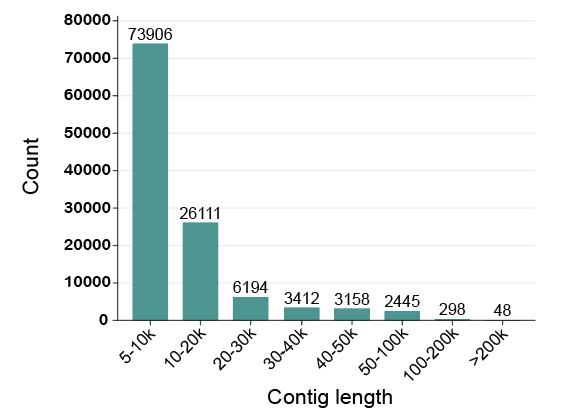

**Figure S8. Length distribution of recovered viral contigs in heavy metal-contaminated soils.**

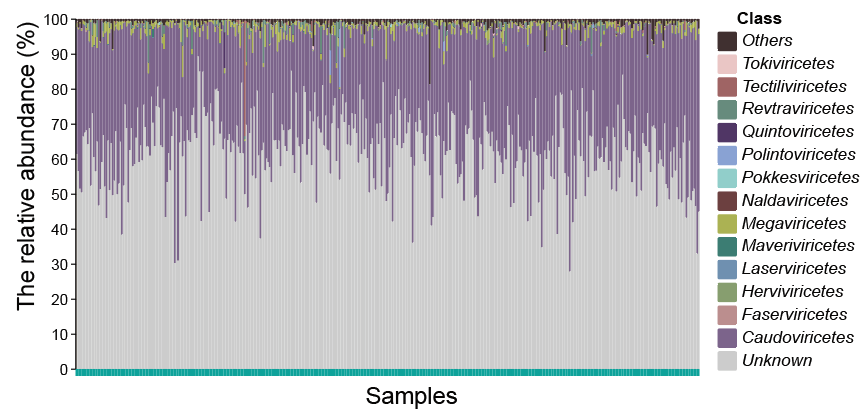

**Figure S9. Taxonomic composition of the viral community of 401 soil metagenomes spanning a gradient of heavy metal contamination across China.**

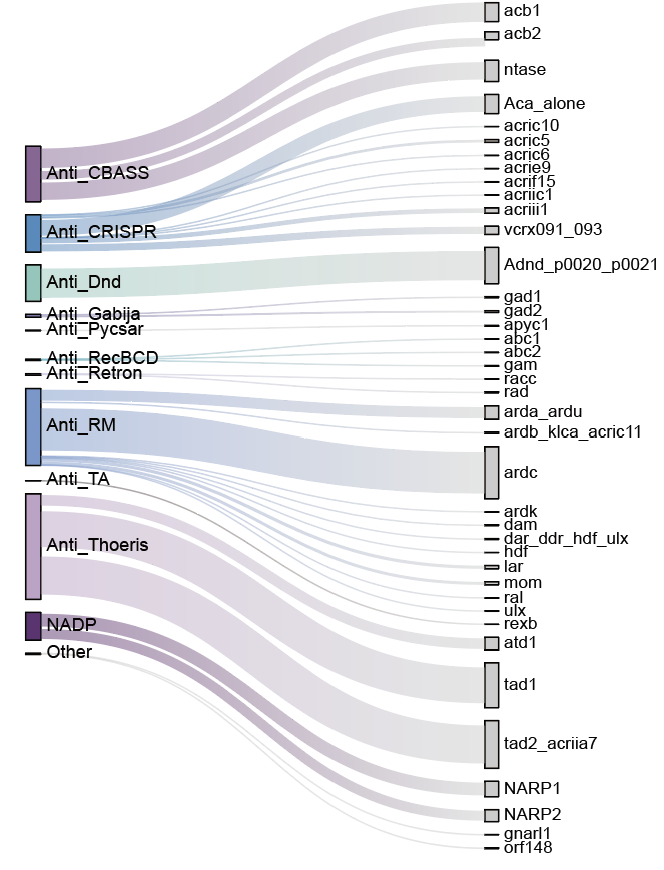

**Figure S10. Subtype composition of ADS encoded by viral communities in 401 heavy metal-contaminated soils.**

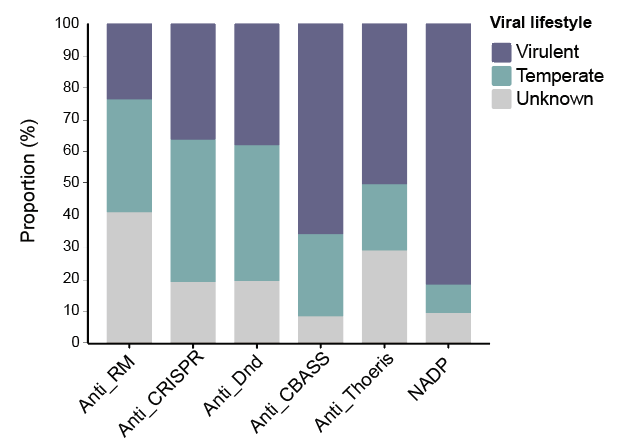

**Figure S11. Distribution of six major ADS across temperate, virulent, and unclassified viruses in heavy metal-contaminated soils.**

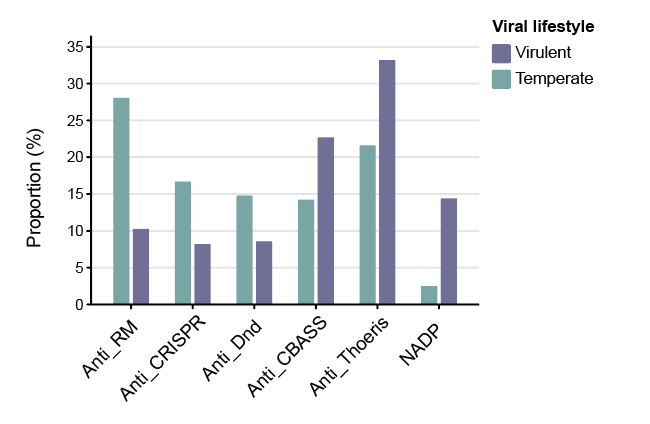

**Figure S12. Relative composition of ADS classes in temperate and virulent viruses from heavy metal-contaminated soils.**

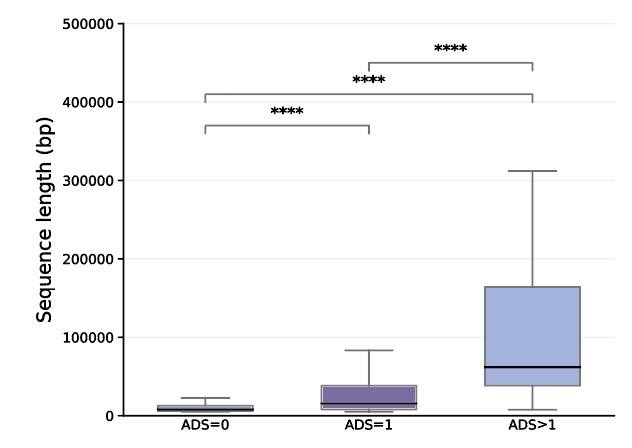

**Figure S13. Genome length distributions of viruses carrying different numbers of ADS in heavy metal-contaminated soils.** (Wilcoxon rank-sum test, ****, *P* < 0.0001)

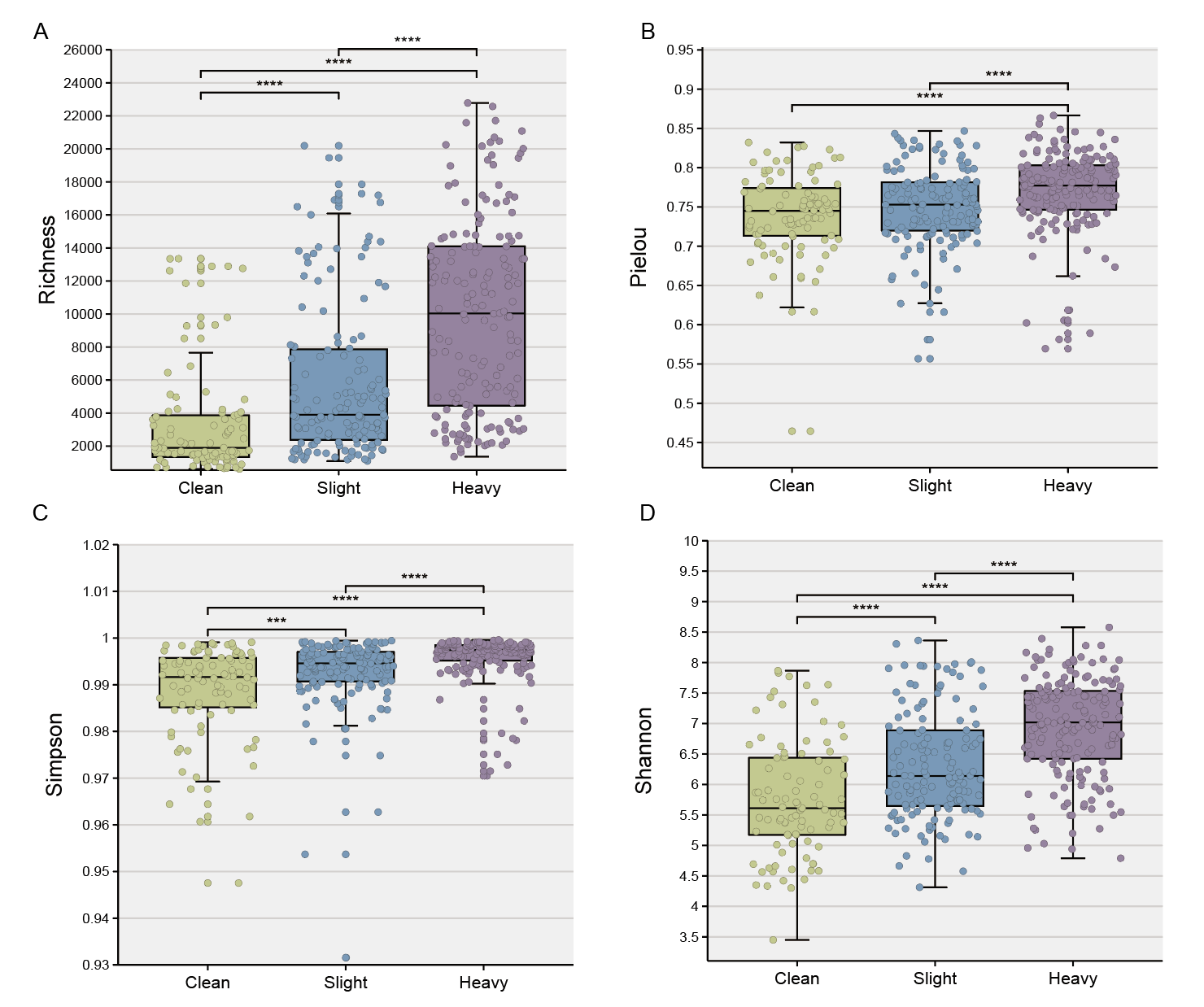

**Figure S14. The alpha diversity of viral community.** (A) Richness index, (B) Pielou index, (C) Simpson, (D) Shannon. (Wilcoxon rank-sum test, ****, *P* < 0.0001)

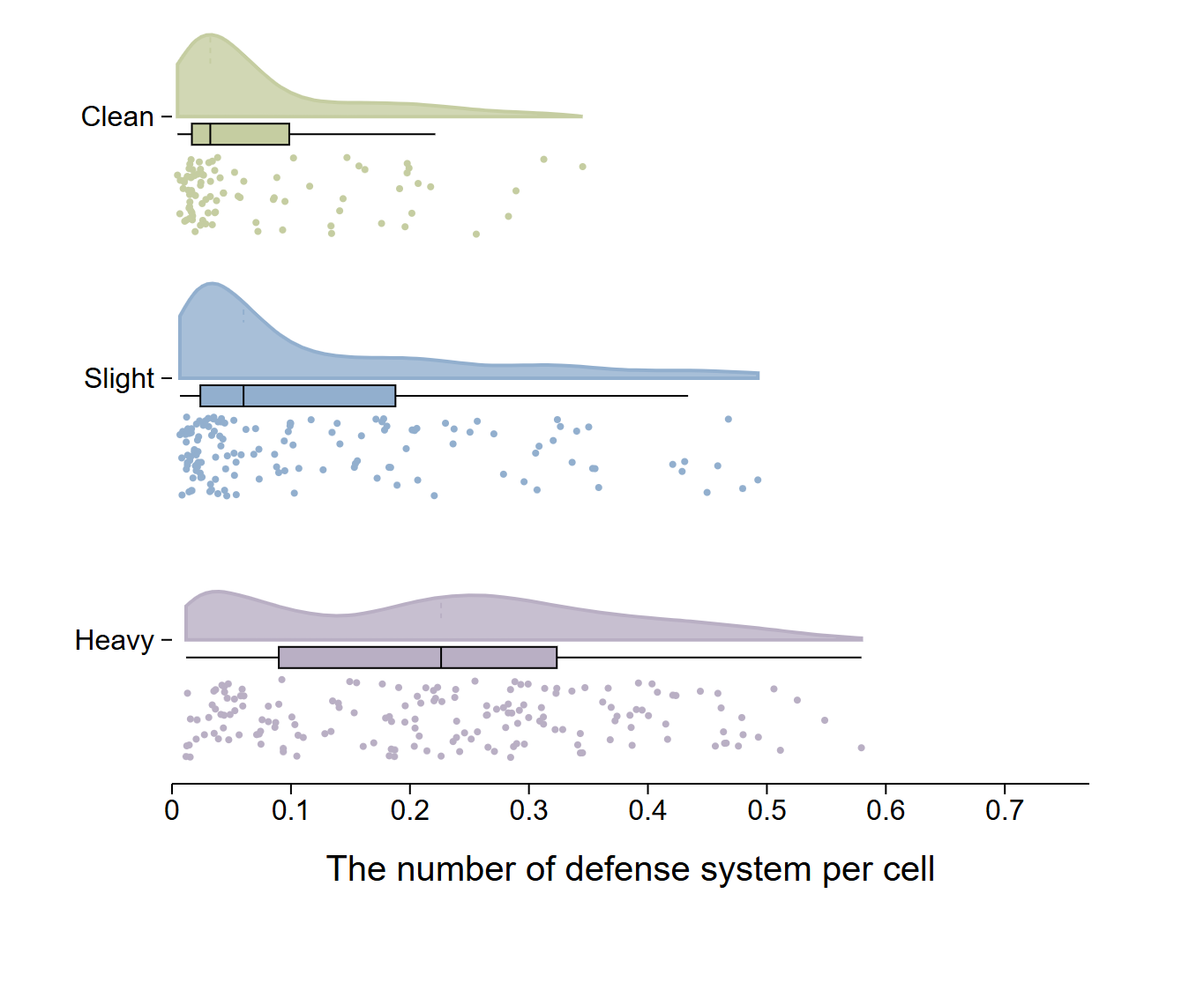

**Figure S15. Variation in the number of antiviral defense genes per prokaryotic genome across the contamination gradient.** Antiviral defense systems were identified from all assembled contigs in each metagenomic sample. The number of detected defense systems was normalized by the estimated number of microbial cell equivalents, which was calculated by ARG-OAP based on 16S rRNA gene counts. The resulting values therefore represent an estimated number of defense systems per cell rather than an absolute count measured for individual cells or reconstructed prokaryotic genomes. Each point represents one metagenomic sample. Density curves show the distribution of values within each contamination category. In the boxplots, the center line represents the median, the box spans the interquartile range (IQR; 25th–75th percentiles), and the whiskers extend to the most extreme values within 1.5 × IQR. Colors denote clean, slightly contaminated, and heavily contaminated soils.

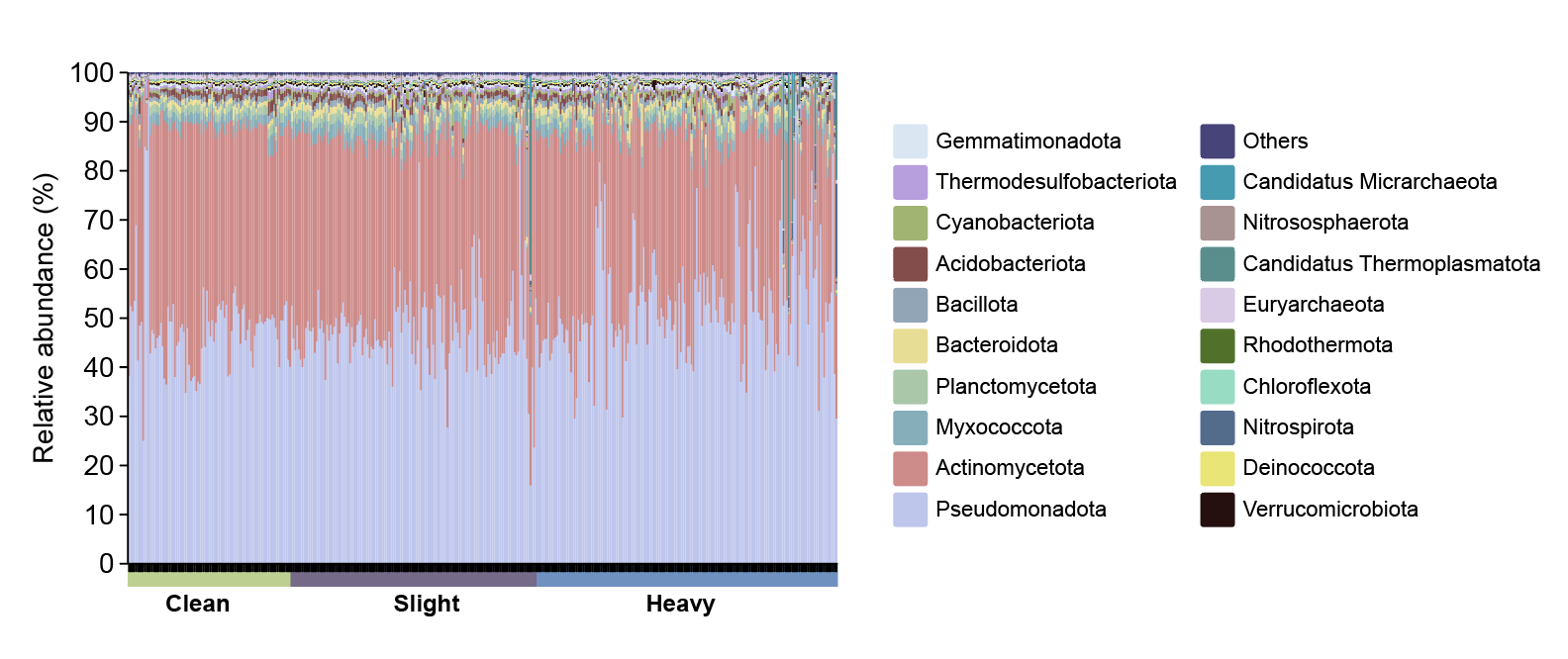

**Figure S16. Phylum-level composition of prokaryotic communities across heavy-metal contamination groups.** Each stacked bar represents one sample (Clean, n = 92; Slight, n = 139; Heavy, n = 170). Colors indicate the relative abundances of bacterial and archaeal phyla based on Kraken2 taxonomic assignments. Phyla not displayed individually are grouped as “Others”.

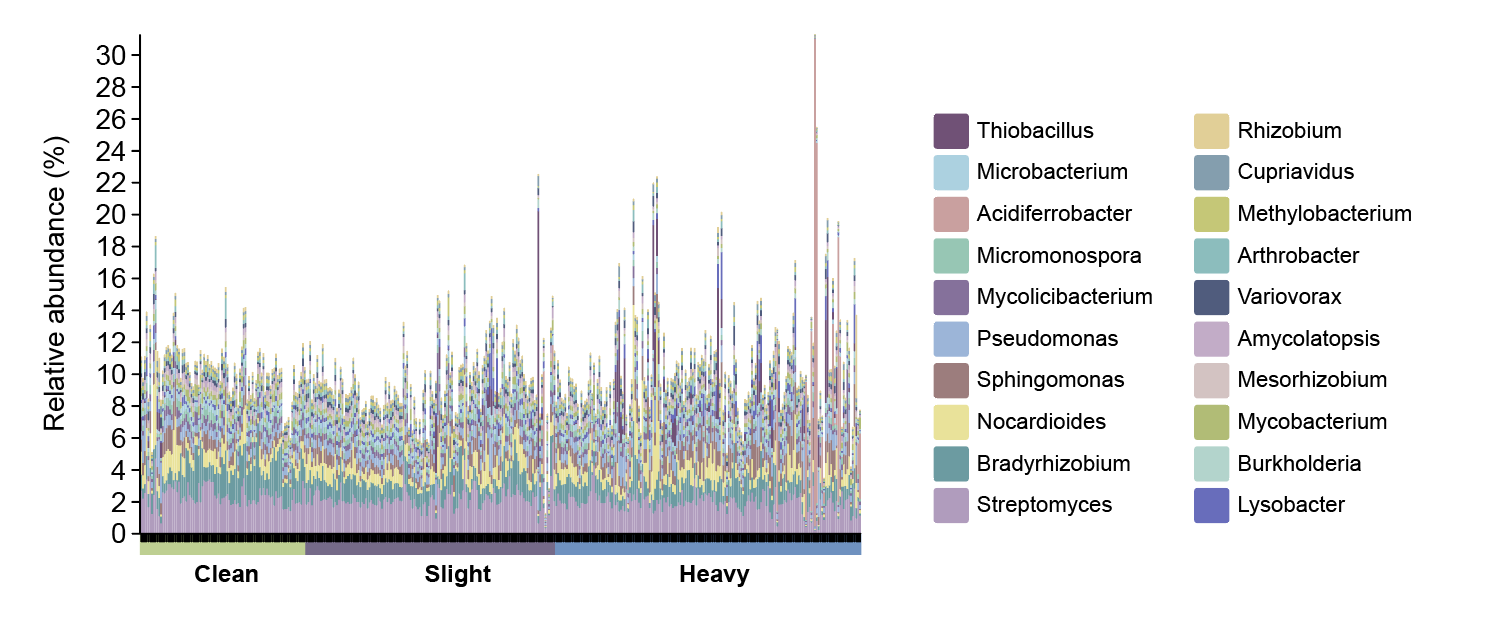

**Figure S17. Relative abundances of the 20 most abundant genera across heavy-metal contamination groups.** Each stacked bar represents one sample, grouped as Clean, Slight or Heavy. The 20 genera were selected by their mean relative abundance across all samples. Other genera and sequences without genus-level assignments are not displayed; therefore, the bars do not sum to 100%.
